# A Bayesian Approach to Restricted Latent Class Models for Scientifically-Structured Clustering of Multivariate Binary Outcomes

**DOI:** 10.1101/400192

**Authors:** Zhenke Wu, Livia Casciola-Rosen, Antony Rosen, Scott L. Zeger

**Affiliations:** Department of Biostatistics and Michigan Institute for Data Science, University of Michigan, Ann Arbor, MI 48109, USA; Division of Rheumatology, Department of Medicine, Johns Hopkins University School of Medicine, Baltimore, Maryland, 21224, USA; Department of Biostatistics, Johns Hopkins University, Baltimore, MD 21205, USA

**Keywords:** Autoimmune disease, Clustering, Dependent Binary Data, Latent Class Models, Markov Chain Monte Carlo, Mixture of Finite Mixture Models

## Abstract

This paper presents a model-based method for clustering multivariate binary observations that incorporates constraints consistent with the scientific context. The approach is motivated by the precision medicine problem of identifying autoimmune disease patient subsets or classes who may require different treatments. We start with a family of restricted latent class models or RLCMs (e.g., Xu and Shang, 2018). However, in the motivating example and many others like it, the unknown number of classes and the definition of classes using binary states are among the targets of inference. We use a Bayesian approach to RCLMs in order to use informative prior assumptions on the number and definitions of latent classes to be consistent with scientific knowledge so that the posterior distribution tends to concentrate on smaller numbers of clusters and sparser binary patterns. The paper derives a posterior sampling algorithm based on Markov chain Monte Carlo with split-merge updates to efficiently explore the space of clustering allocations. Through simulations under the assumed model and realistic deviations from it, we demonstrate greater interpretability of results and superior finite-sample clustering performance for our method compared to common alternatives. The methods are illustrated with an analysis of protein data to detect clusters representing autoantibody classes among scleroderma patients.

## 1. Introduction

Autoantibodies are the immune system’s response to specific cellular protein complexes or “machines” (e.g., Rosen and Casciola-Rosen, 2016). Differential immune responses mounted towards the machines may lead to strikingly different clinical trajectories between autoimmune disease patients. Using autoantibodies as probes, a precision medicine goal is to discover novel autoimmune disease patient subsets that require different treatments prior to major clinical manifestation. This paper is motivated by the need to first formulate a model that respects the scientific constraint that the immune system responds to machines, that is to all components, rather than to individual proteins, and second, based on the model, to perform clustering in a Bayesian framework using autoantibody test data.

To illustrate the scientific constraint, Figure 1 shows a hypothetical patient with binary state vector ***η***_*i*_ = (1, 0, 1)^T^ in the middle panel, indicating that her immune system produced autoantibodies to the proteins (autoantigens) in Machines 1 and 3 but not Machine 2. The right panel of Figure 1 shows an illustrative example of three different machines (*M* = 3) with orthogonal machine profiles, though the proposed approach will cover general non-orthogonal cases. The binary matrix *Q* specifies which proteins constitute each machine. We refer to the rows of *Q* as “machine profiles” where *Q*_*m𝓁*_ = 1 if protein *𝓁* is a component of machine *m*. The left panel shows Γ_*i𝓁*_, which represents the error-free true presence or absence of autoantibody against protein *𝓁* = 1, …, *L* for subject *i* = 1, …, *N*. The entire row Γ_*i∗*_ = (Γ_*i*1_, …, Γ_*iL*_) is highlighted for subject *i*. The multivariate binary data for subject *i* (***Y***_*i*_, not shown in the figure) are Γ_*i∗*_ measured with error. The statistical goal is to incorporate the scientific constraint when clustering patients into subsets with distinct responses to the machines. The choice of binary rather than continuous measures of autoantibodies is specific to the gel electrophoresis and autoradiogram (GEA) technology that produced the data where the amount of protein is not informative about clustering patients (Wu et al., 2019).

**Figure 1:**
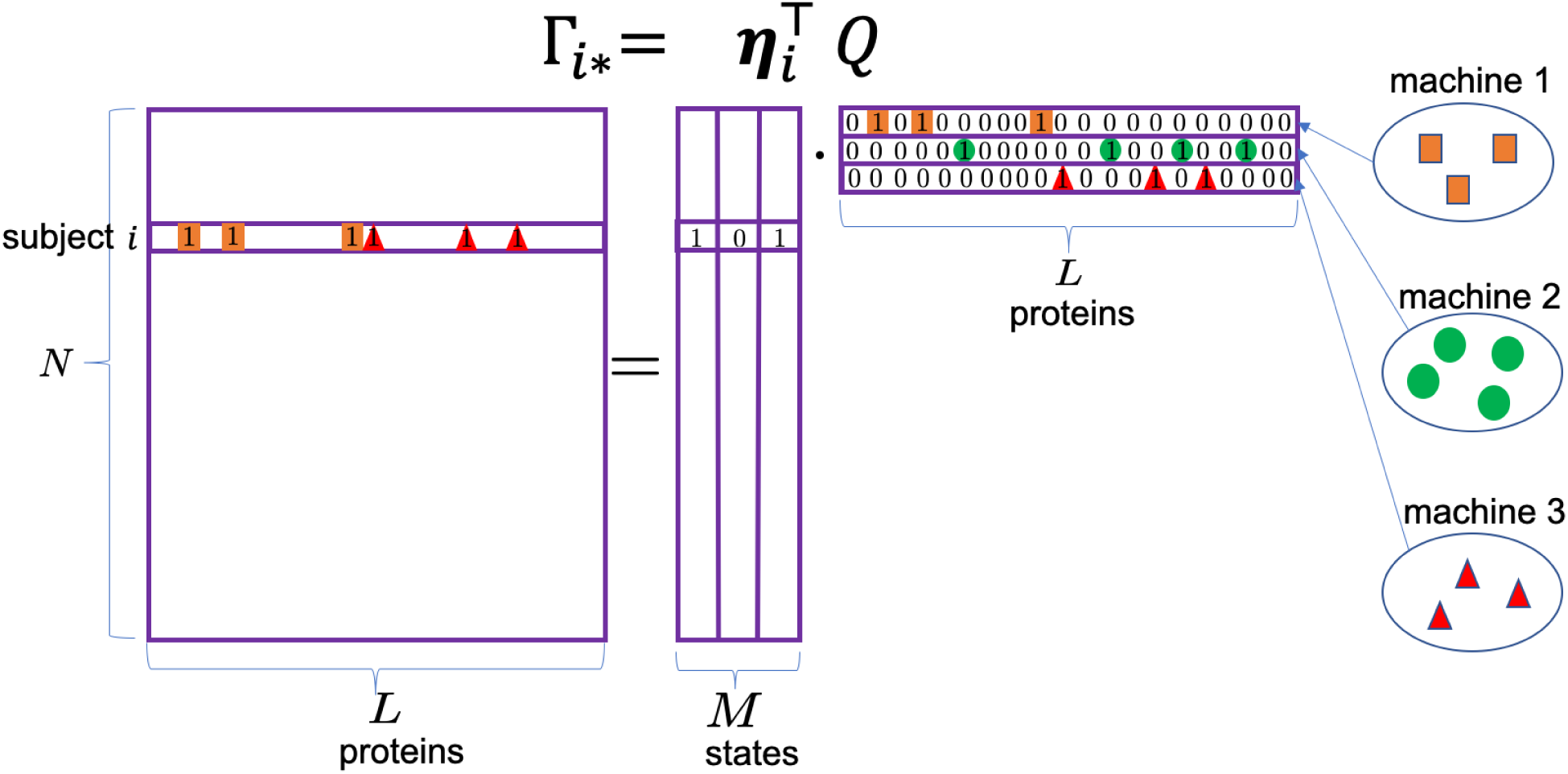
The scientific constraint means that immune responses are mounted towards all protein components in each machine rather than towards individual proteins. For illustrative purposes, the machine profiles here are orthogonal so the true presence or absence of proteins can be represented by binary matrix factorization; the proposed approach covers general non-orthogonal cases. (This figure appears in color in the electronic version of this article, and any mention of color refers to that version.)

The knowledge that the immune system attacks machines of multiple proteins rather than single proteins is why we refer to this approach as scientifically-structured clustering (SSC). We use “scientific structure” to refer to latent binary variables that may be interpreted as having a scientific meaning. For example, in the protein data example, each element of the latent binary random vector ***η***_*i*_ represents the presence or absence of a machine. The scientific meaning of a machine is defined by its component proteins indicated by the corresponding row of *Q*. SSC for multivariate binary data has a number of potential applications beyond the example here. For example, in cognitive diagnosis, the latent states represent basic cognitive abilities and the 1s in *Q* represent the test items requiring each ability (e.g., Junker and Sijtsma, 2001); in epidemiology, the latent states can indicate unobserved disease-causing agents and the 1s in *Q* represent the molecular targets tied to each causative agent (Wu et al., 2016, e.g.,). Most importantly, in all these applications, the resulting clusters conform to the existing scientific context and therefore can be used to address relevant questions.

As detailed below, the model proposed to address the motivating patient clustering problem is a member of the family of restricted latent class models or RLCMs (e.g., Xu and Shang, 2018). In our motivating example, however, the possible combinations of machines that the immune systems can target in the population of patients are often unknown. This means not knowing the set of distinct population-level latent state patterns *𝒜*, neither the size |*𝒜*| nor its elements. In addition, the definitions of machine profiles *Q* are often unknown.

In addressing the motivating problem, this paper makes a primary contribution to the literature on RLCMs. In a Bayesian framework, we allow an unknown number of classes in a mixture of finite mixture (MFM, Miller and Harrison, 2018) framework where each component has multivariate discrete parameters, e.g., the binary states in our context. We then derive a posterior sampling algorithm based on Markov chain Monte Carlo (MCMC) featuring split-merge updates that directly and efficiently sample the clustering allocations. The algorithm conveniently overcomes the hard constraint |*𝒜*| 2^*M*^ by post-processing to merge sampled clusters having the same sampled state vectors. The model works with an unknown number of classes, unknown set of state patterns *𝒜*, and unknown *Q*-matrix. We use informative prior assumptions on the number and definition of latent classes to be consistent with the scientific knowledge, so that the posterior distribution tends to concentrate on fewer clusters with sparser latent state patterns.

The rest of the paper is organized as follows: Section 2 specifies the proposed model and prior distribution. Section 3 derives the MCMC algorithm for posterior inference. Section 4.1 presents simulation studies to show the better clustering performance of the proposed clustering method when compared to three common alternatives. Section 4.2 illustrates the methods with an analysis of protein data for subsetting scleroderma patients. The paper concludes with a discussion of model extensions and limitations.

## 2. Models

First formulated by Lazarsfeld (1950), latent class models (LCMs) have become an important tool for modeling multivariate discrete responses (e.g., Goodman, 1974; Dunson and Xing, 2009) and model-based clustering (e.g., Vermunt and Magidson, 2002). Alternative parametric approaches focus on underlying continuous variable specifications (e.g., Albert and Chib, 1993) which incorporate latent multivariate Gaussian random variables that are linked to the categorical observations through thresholding. However, these models do not directly perform clustering. In this paper, we focus on extensions of the classical LCMs to incorporate scientific constraints when clustering multivariate binary responses.

*Notation*. Let ***Y***_*i*_ = (*Y*_*i*1_, …, *Y*_*i𝓁*_)^T^ be the vector of binary measurements for subject *i* = 1, …, *N* and feature *𝓁* = 1, …, *L*. Let **Y** collect all the observations into an *N* × *L* binary data matrix. Let Γ_*i𝓁*_ indicate the true presence/absence of feature *𝓁* for observation *i*. Let Γ be an *N* × *L* binary matrix with (*i, 𝓁*)-th element being Γ_*i𝓁*_, referred to as design matrix. We use Γ_*i∗*_ = (Γ_*i*1_, …, Γ_*iL*_) and Γ_*∗𝓁*_ = (Γ_1*𝓁*_, …, Γ_*N𝓁*_)^T^ to represent the *i*-th row and *𝓁*-th column, respectively. Let ***η***_*i*_ = (*η*_*i*1_, …, *η*_*iM*_)^T^ represent the unobserved latent states for subject *i* = 1, …, *N*. Let ***η***_*i*_ lie in an unknown population state space *𝒜* ⊆ {0, 1}^*M*^. We use |*𝒜*| to denote the cardinality of *𝒜*. As a result, there are |*𝒜*| distinct patterns of latent states. We refer to *𝒜* = {0, 1}^*M*^ that contains all binary patterns as “saturated” and otherwise “unsaturated”. We collect the latent states into an *N* × *M* matrix *H* with (*i, m*)-th entry being *η*_*im*_. Let *Q* be an unknown *M* × *L* binary matrix with each row specifying the scientific meaning of each latent state. Let *Q*_*m∗*_ = (*Q*_*m*1_, …, *Q*_*mL*_) and *Q*_*∗𝓁*_ = (*Q*_1*𝓁*_, …, *Q*_*M𝓁*_)^T^ represent row *m* and column *𝓁* of *Q*, respectively. Let [*A* | *B*] represent the conditional distribution of random quantity *A* given another random quantity *B*. When *B* = ∅, [*A*] represents the distribution of *A*. Let I(*A*) be an indicator function which equals 1 if statement *A* is true and 0 otherwise.

### 2.1 Proposed Model

We specify the model for the motivating example in two steps: i) impose scientific structure upon the actual presence or absence of proteins (Γ_*i∗*_), and ii) parameterize [***Y***_*i*_ | Γ_*i∗*_]. The first step is needed to respect existing biological knowledge in the scientific context and the second step characterizes the measurement process.

Let the true presence or absence of feature *𝓁* for subject *i* = 1, … *N* be

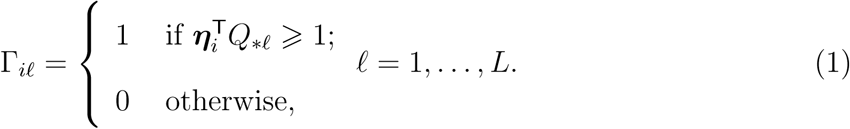

To emphasize the dependence of Γ_*i𝓁*_ on *Q* and ***η***_*i*_ for *𝓁* = 1, …, *L*, we write the row vector Γ_*i∗*_ = Γ(*Q*, ***η***_*i*_) so that Γ(*Q*, ***η***_*i*_) _*𝓁*_ = Γ_*i𝓁*_. In addition, let Γ(*Q, 𝒜*) represent an |*𝒜*| × *L* binary matrix with each row representing the true presence or absence of feature *𝓁* = 1, …, *L* for a subject having a particular latent state pattern in *𝒜*.

Next we specify the measurement error model:

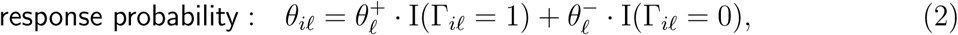

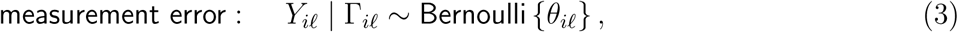

independently for feature *𝓁* = 1, …, *L* and subject *i* = 1, …, *N*. Borrowing the terminologies in the statistical literature for diagnostic testing, we characterize the stochastic discrepancies between the actual Γ_*i𝓁*_ and observed presence/absence of autoantibodies by sensitivity 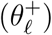 and specificity 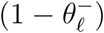. Let 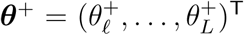 and 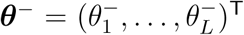. In this paper, we assume 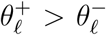, because the diagnostic test based on immunoprecipitation (IP) is known to be both sensitive and specific. The sensitivity for a given protein is assumed to be the same regardless from which machine(s) it comes. Importantly, both the sensitivities and specificities can vary across proteins.

Equations (1) to (3) assume the probability of observing a protein given that it is present does not depend on how many machines it is present in, that is, it depends only on I 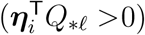 but otherwise not on the value of 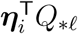. In our application, the choice is driven by the technology used. The ability to detect the protein does not depend on how many machines produced it. In addition, the amount is irrelevant for clustering patients. Further the model assumes binary response/nonresponse to each protein, rather than quantifying the response as a real value. This modeling choice follows the convention that clinicians interpret the presence or absence of bands corresponding to each protein on the images obtained from GEA (Wu et al., 2019).

Equations (1) to (3) are related to some existing models proposed in cognitive diagnosis (e.g., Templin and Henson, 2006, known *Q*) and epidemiology (e.g., Wu et al., 2016, *Q* = *I*_*L*×*L*_), which are examples of restricted LCMs (RLCM, Xu, 2017). In addition, imposing symmetric error rates 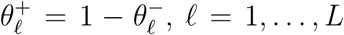, results in the one-layer model of Rukat et al. (2017). The model can also be viewed as Boolean matrix factorization (BMF, Miettinen et al., 2008) because model (1) is equivalent to 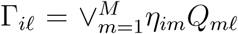 where the logical “OR” operator “V “outputs one if any argument equals one. The rows in *Q* are basis patterns for compactly encoding the *L* dimensional Γ_*i∗*_ vector by *M* (≪*L*) bits in ***η***_*i*_. When *Q* has orthogonal rows, BMF further reduces to nonnegative matrix factorization Γ = *HQ* (NMF, Lee and Seung, 1999). Zhang et al. (2007) proposed optimization methods to obtain the best factorization while Meeds et al. (2007) and Ni et al. (2019) took a nonparametric Bayes approach. In contrast to these work, we add a layer of mixture model in (5) to (10) that provides additional parsimony and posterior inference of the number of classes.

### 2.2 Identifiability Considerations for Posterior Algorithm Design

In general, identifiability conditions based on model likelihood can guide the design of simulation studies by informing the choice of the simulation truths that are statistically identifiable. In real data analysis, although truth is not expected to be known even if the model is a reasonable approximation, identifiability conditions can still dramatically reduce the size of parameter space.

Specifically, under a fixed *M* and possibly unsaturated *𝒜*, Gu and Xu (2019a) provided sufficient conditions on Γ(*Q, 𝒜*) for strictly identifying *𝒜*, ***π, θ***^+^, ***θ***^−^ and Γ(*Q, 𝒜*). Although Γ(*Q, 𝒜*) is identifiable, *Q* may only be identified up to its equivalence classes, where *Q*_1_ and *Q*_2_ are equivalent if Γ(*Q*_1_, *𝒜*) = Γ(*Q*_2_, *𝒜*) with equality holds elementwise.

In our simulations and data application, we restrict the inference algorithm to be performed on the set of *M* × *L* binary *Q*-matrices:

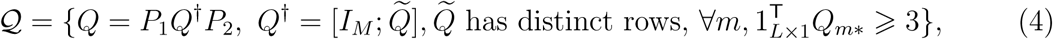

where *P*_1_ and *P*_2_ are *M* - and *L*-dimensional permutation matrices. First, under a saturated *𝒜*, the conditions in (4) are based only on *Q*, easy-to-check, and necessary and sufficient for statistical identification of ***π, θ***^+^, ***θ***^−^ and *Q* (Gu and Xu, 2019b). Second, the constraint *𝒬* also greatly facilitates posterior sampling by focusing on a small subset of binary matrices. In fact, among all *M* by *L* binary matrices, the fraction of *Q* ∈ *𝒬* is at most 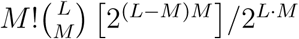 and quickly decays as the number of machines *M* increases.

We now turn to inferring individual latent states based on complete-data likelihood [**Y** | *H, Q*, ***θ***^+^, ***θ***^−^]. Even given *Q*, conditions for identifying *H* exist but may fall short of ensuring consistent estimation of *H* because the number of unknowns in *H* diverges as the sample size increases. Consistent estimation requires extra conditions, e.g., the number of measurements *L* increases with the sample size (Chiu et al., 2009). In finite samples and dimensions, we address this issue in a Bayesian framework by encouraging *H* to be *a priori* of low complexity, i.e., few classes of distinct and sparse latent state vectors.

Finally, in real data analysis, incorporating informative priors, e.g., partially-known *Q* or high sensitivity and specificity may alleviate theoretical identifiability issues (e.g., Wu et al., 2016). The prior uncertainty for the non-identified parameters will propagate into the posterior and not vanish even as the sample size approaches infinity (e.g., Kadane, 1975).

### 2.3 Priors

We first specify a prior for allocating observations into clusters and then a prior for the multivariate binary state vectors in the clusters.

#### Prior with an unknown number of classes

Though used interchangeably by many authors, we first make a distinction between a “component” that represents one of the true mixture components in the specification of a mixture model (referred to as “classes” in LCMs and this paper) and a “cluster” that represents a block of observations grouped together. Let *K* be the number of mixture components in the population and *T* the number of clusters in the sample (Miller and Harrison, 2018). We have *T* ⩽ *K* which means there exist *T* – *K* empty components not realized in a random sample.

Additional notation is needed for our prior specification. Let *Z*_*i*_ ∈ {1, 2, …, *K*} be the component indicators, and let ***Z*** = {*Z*_*i*_, *i* = 1, …, *N*}. Let *C*_*k*_ = {*i* : *Z*_*i*_ = *k*} be the subjects in component *k*, and *C* = {*C*_*k*_ : |*C*_*k*_| *>* 0, *k* = 1, …, *T*} be *T* observed clusters. Note the partition *C* is invariant to component relabeling. Let *C*_−*i*_ = {*C*_*k*_ *\* {*i*} : |*C*_*k*_ *\* {*i*}| *>* 0} be the clusters excluding subject *i*. Let **Y**_*C*_ = {***Y***_*i*_, *i* ∈ *C*} be the data in cluster *C* ∈ *C*. Finally let ***α***_*k*_ ∈ {0, 1}^*M*^ be the latent state vectors for component *k* = 1, …, *K*.

We specify a prior distribution of clustering allocations *C* as follows:

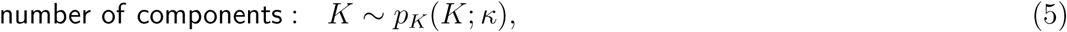

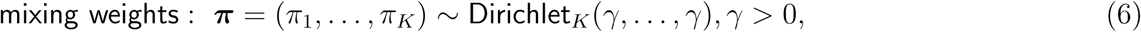

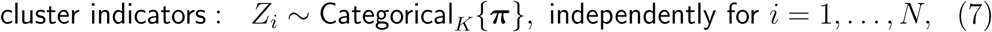

where *p*_*K*_(*K*; *κ*) is a distribution over *K* = 1, 2, …, e.g., Geometric(*κ*). The prior for *𝒞* is

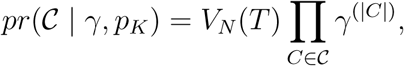

where 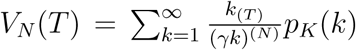, and *k*^(*n*)^ = *k* · (*k* + 1) *…* (*k* + *n* − 1), *k*_(*n*)_ = *k* · (*k* − 1) *…* (*k* − *n* + 1), and *k*^(0)^ = *k*_(0)_ = 1, *k*_(*n*)_ = 0 if *k < n* (Miller and Harrison, 2018).

*Prior for* ***α***_*k*_ *in each component*. We specify a prior distribution for the component-specific parameters ***α***_*k*_ to encourage sparser binary patterns:

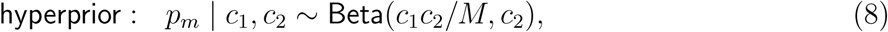

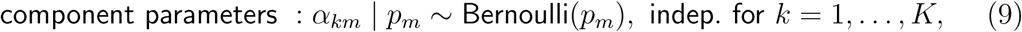

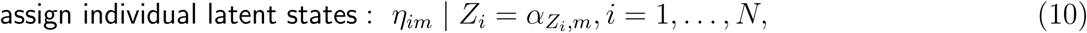

independently for *m* = 1, …, *M*. Equations (8) to (10) induce a marginal prior [***α***_1_, …, ***α***_*K*_ | *c*_1_, *c*_2_] upon integrating over ***p*** = (*p*_1_, …, *p*_*m*_)^T^ (Web Appendix A1.1) and is a truncated Indian Buffet Process (Ghahramani and Griffiths, 2006). In what follows, we set *κ* = 0.1 and *c*_2_ = 1 which offer good clustering results in simulations and data analysis. We reparametrize *c*_1_ in terms of 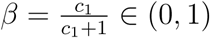 and specify the prior *β* ∼ Beta(*a*_*β*_, *b*_*β*_) where *a*_*β*_ = *b*_*β*_ = 1.

##### Remark 1

A reviewer raised a question about model specification that, given a finite *M*, by following (8) to (9) to sample {*η*_*im*_, *i* = 1, …, *N*} instead of *α*_*km*_’s, there is already a positive prior mass on the equality of two binary vectors with finite length 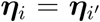 (hence *i* and *i*′ belong to the same cluster), why is it necessary to have an additional layer of clustering structure through mixture models (5) to (7)? In this paper, we choose the mixture model specification because it has a prior on the number of classes which allows additional parsimony by *a priori* encouraging few classes and posterior inference on the number of classes. Although |*𝒜*| ⩽ 2^*M*^, *K* in the prior is not upper bounded. With an unbounded *K*, we can build on the algorithm of Miller and Harrison (2018) for inferring the number of classes. Because the component parameters are multivariate binary, we do need an additional step of merging clusters with the same latent states via *post hoc* processing at each MCMC iteration. The algorithm with merging is applicable to general MFM models with discrete component parameters. Web Appendix A1.2 discusses the merge operation.

For the rest of parameters, let the prior for *Q* be the uniform distribution over *𝒬* in (4). For ***θ***^+^ and ***θ***^−^, we specify 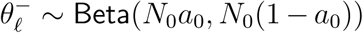, 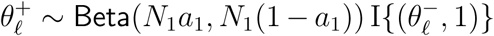, independently for *𝓁* = 1, …, *L*. Taken together, the joint distribution is

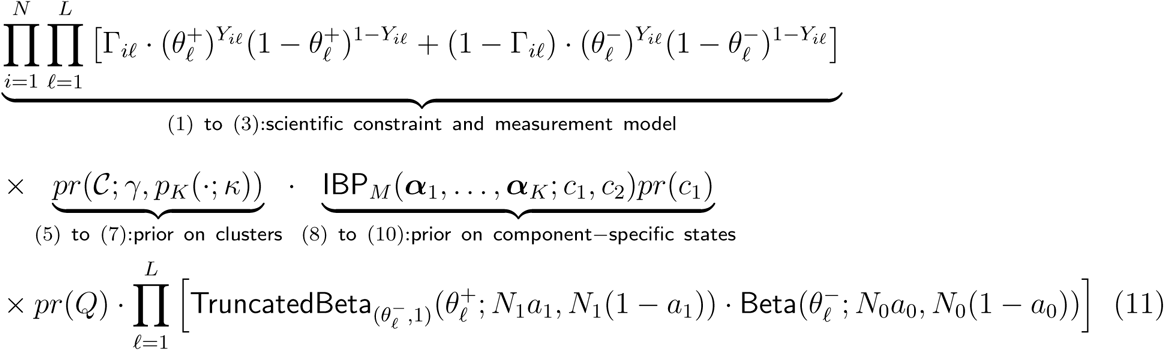

Figure 2 shows a schematic representation of the hierarchical model via directed acyclic graph (DAG) describing the relationships between the parameters and observations.

**Figure 2:**
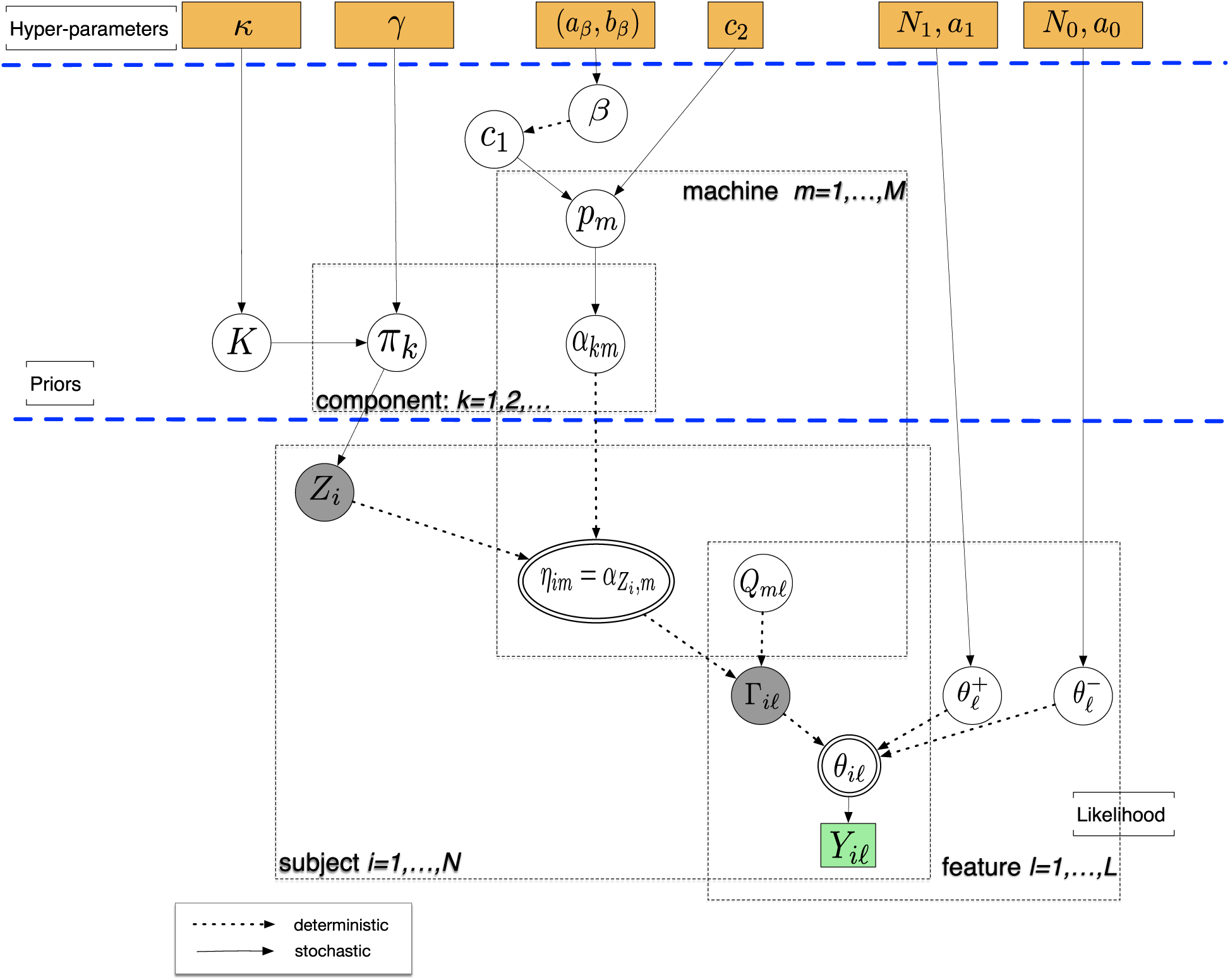
The directed acyclic graph representing the model structure. The quantities in squares are either data or hyperparameters; the unknown quantities are shown in the circles. The double-stroke circle represents multiplexer variable, which copies the value of one of its parent nodes chosen by the selector variable (shaded). The arrows connecting variables indicate that the parent parameterizes the distribution of the child node (solid lines) or completely determines the value of the child node (dotted arrows). The rectangular “plates” where the variables are enclosed indicate that a similar graphical structure is repesated over the index; the index in a plate indicates subjects, clusters, machines, or features. (This figure appears in color in the electronic version of this article, and any mention of color refers to that version.)

## 3. Posterior Inference

We develop inferential procedures to address the following three questions: 1) how many scientific clusters are in the data; 2) what are the unique latent states present in the sample, and 3) what is the latent state vector for every observation. To approximate the posterior distribution we use an MCMC algorithm for the inference of any functions of unknown parameters and latent variables (Gelfand and Smith, 1990). We focus on presenting the posterior sampling algorithm with a finite *M* which is effective in our application by treating *M* as an upper bound, though the posterior algorithm for infinite *M* is readily derived by following Teh et al. (2007).

### 3.1 MCMC Algorithm

When the number of components *K* is unknown, one class of techniques updates component-specific parameters along with *K*. For example, the reversible-jump MCMC (RJ-MCMC, Green, 1995) works by an update to *K* along with proposed updates to the model parameters which together are then accepted or rejected. However, designing good proposals for high-dimensional component parameters can be non-trivial. Alternative approaches include direct sampling of *K* (e.g., Nobile and Fearnside, 2007; McCullagh et al., 2008). Here we build on the algorithm of Miller and Harrison (2018) for sampling clusters with a prior on the number of mixture components.

Step 0: Initialize all parameters from their priors except for *Q* and clustering allocations *C*. For *Q*, we set all elements to be zero except for columns *𝓁* that satisfy *N* ^−1^ Σ*_i_ Y*_*i𝓁*_ *> τ*_1_, for which we initialize by 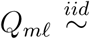 Bernoulli(*p*), *m* = 1, …, *M*, where *p* and *τ*_1_ are pre-specified. We set *p* = 0.1 and *τ*_1_ = 0.3 which work well in our simulations and data analysis. For *C*, we initialize with one cluster comprised of all subjects.

For iteration *b* = 1, …, *B*, repeat Steps 1-7:

Step 1: Update *C* by directly sampling from its posterior without the need for considering component parameters or empty components. For subject *i* = 1, …, *N*, we assign subject *i* to an existing cluster *C* ∈ *𝒞*_−*i*_ or a new one on its own with probabilities:

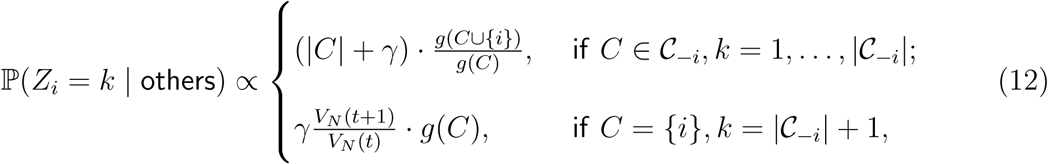

where *g*(*C*) is the marginal likelihood for data **Y**_*C*_ and is computed in Web Appendix A2. The update (12) favors forming a cluster of its own {*i*} if adding the *i*-th observation to any existing cluster fits poorly with data in that cluster. Following the split-merge recipe in Jain and Neal (2004) that efficiently explores the large space of clustering allocations (see details in Web Appendix A3), we build on (12) to update *C* globally which creates or removes clusters for multiple subjects at a time.

Step 2: Update ***α***_*k*_ by the distribution over {0, 1}^*M*^ :

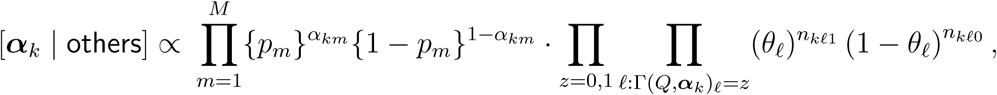

for cluster *k = 1;* …; *T*, where 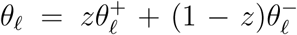, and 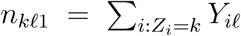 and 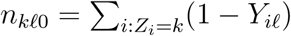. Set ***η***_*i*_ = ***α***_*Z*_*i*, for subject *i* = 1, …, *N*.

At the end of iterations, if some of the discrete mixture component parameters {***α***_*k*_, *j* = 1, …, *T*} are duplicated, we post-process the posterior samples by merging clusters in *𝒞* associated with identical latent states to obtain scientific clusters with distinct latent states; denote scientific clusters by 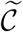 and 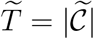.

Step 3: Update the measurement error parameters for *𝓁* = 1, …, *L*:

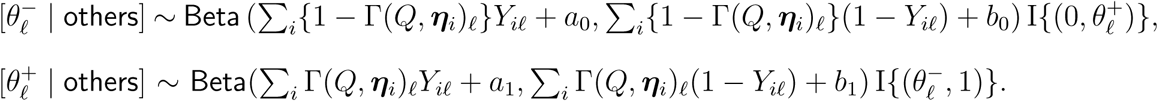

Step 4: Update hyperparameter *c*_1_ = *β/*(1 − *β*) by updating *β*:

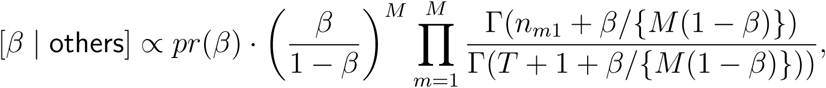

where 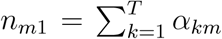 is the number of observed clusters with activated *m*-th state. The update can be done based on a dense grid over (0, 1) (Hoff, 2005).

Step 5: Update *p*_*m*_ ∼ Beta(*n*_*m*1_ + *c*_1_*/M, T* − *n*_*m*1_ + 1), independently for *m* = 1, …, *M*.

Step 6: Update *Q* via constrained Gibbs sampler. For *𝓁* = 1, 2, …, *L, m* = 1, 2, …, *M*, do not update *Q*_*m𝓁*_ if i) *Q*_*∗𝓁*_ = ***e***_*m*_ where ***e***_*m*_ is a column vector of with the *m*-th element being 1 and 0 for other elements, ii) 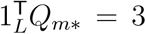 and *Q*_*m𝓁*_ = 1, or iii) 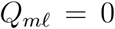 in a column of 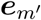 with *m*′ ≠ *m*, and there is a single 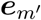 in the columns of the current *Q*. Otherwise, we follow an efficient update in Liu (1996) by flipping *Q*_*m𝓁*_ from 1 − *z* to *z* with probability ℙ(*Q*_*m𝓁*_ = *z* | others)*/*{1 − ℙ (*Q*_*m𝓁*_ = *z* | others)}, where

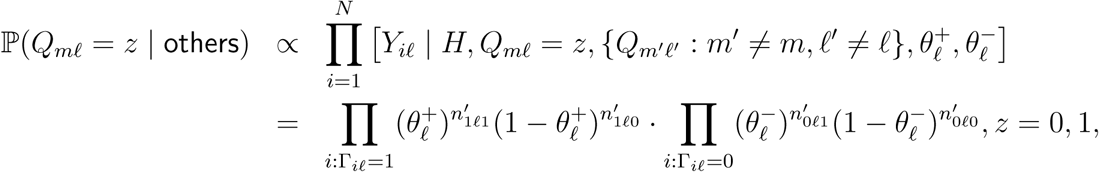

where 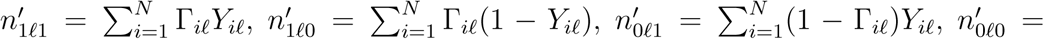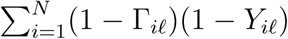. In this step, we also identify “partner latent states” and merge the corresponding rows in *Q*. Specifically, we collapse two states (*m, m*′) that are present or absent together among subjects 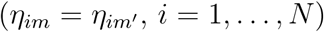. We set 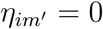 for any *i* and 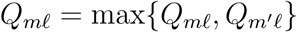, *𝓁* = 1, …, *L*. It is easy to verify that this scheme preserves conditions (4) for *Q* truncated to the rows corresponding to active states. The merging scheme readily generalizes to cases with more than two partner states. Finally, we reset to zeros for the rows of *Q* corresponding to the unused latent states at an iteration.

We have used a few practical strategies to improve the sampling from many discrete parameters, e.g., {***α***_*k*_, *k* = 1, …, *T*} and *Q*. First, the prior that encourages fewer clusters propagates into the posterior so large *T* s are visited less frequently. Second, we put an upper bound on *M* in real applications followed by sensitivity analysis. Third, *Q* is restricted to lie in a subset of {0, 1}^*M* ×*L*^ informed by condition (4). Finally, in our experience, more efficient exploration of clustering allocations among the observations by global split-merge updates helps the sampling of *Q*. Convergence checks are presented in Web Appendix A4.

### 3.2 Posterior Summaries

Let 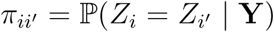 be the posterior co-clustering probabilities for any two subjects *i* and *i*′. We estimate 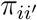 by the empirical frequencies 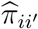 of subjects *i* and *i*′ being clustered together across MCMC iterations. For point estimation, we compute the least square (LS) clustering 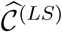 by minimizing the squared distance to the 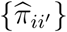, arg min_*b*_ 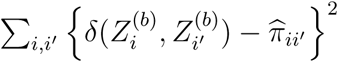, where *δ*(*a, a*′) = 1 if *a* = *a*′ and zero otherwise (Dahl, 2006). In addition, because the interpretation of ***η***_*i*_ depends on *Q*, we choose an optimal *Q* from the posterior samples that minimizes a loss function. We select an iteration *b*^*∗*^ that minimizes the loss: *b*^*∗*^ = arg min 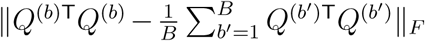 where 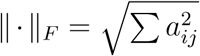 is matrix Frobenius norm. We denote it by 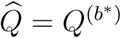. Turning to the inference of *H* = {***η***_*i*_}, we rerun the algorithm by fixing 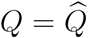 which produces additional posterior samples to reduce the Monte Carlo error in approximating 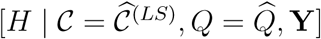.

## 4. Results

We illustrate the utility of Bayesian RLCM on both simulated and real data. First, we assess the performance of Bayesian RLCM on cluster estimation under simulation scenarios corresponding to varying levels of measurement error, dimension, sparsity level of each machine, sample size and mixing weight. Using data simulated under the assumed RLCM and realistic deviations from it, the proposed inference algorithm performs clustering as well as or better than common alternative binary-data clustering methods. We first analyze a single randomly generated data set to highlight differences among the methods. We then use independent replications to evaluate frequentist performance of Bayesian RLCM in cluster estimation and contrast with the alternatives. Web Appendix A5.1 details the simulation scenarios. To assess the chance-corrected agreement between the true and estimated clustering allocations of the same observations, we used the adjusted Rand index (aRI, Hubert and Arabie, 1985).

See Web Appendix A5.2 for details. The value of aRI is between −1 and 1 with values closer to 1 indicating better agreement. Finally, data from scleroderma patients are analyzed.

### 4.1 Simulated Examples to Study Model Performance

#### Simulation 1: More accurate clustering through scientifically structured classes

Figure 3 shows a random data set **Y**, the design matrix Γ, as well as the clusters obtained using complete-linkage, Hamming distance hierarchical clustering (HC), standard eight-class Bayesian latent class analysis (LCA, e.g., Garrett and Zeger (2000)), subset clustering analysis (SC; Hoff, 2005) and our Bayesian RLCM with unknown number of clusters fitted with truncation level *M* ^*†*^ = 5. In this setting, HC is sensitive to noise and tends to split a true cluster or group observations from different true clusters. Unlike the others, the Bayesian RLCM automatically selects and filters subsets of features that distinguish eight classes (through scientific structures in (1)) and hence has superior clustering performance producing clusters that agrees quite well with the truth. For illustrative purposes, we showed an extreme example to highlight the different performances on a single random data set. Although the proposed model-based approach does not always perfectly reconstruct the clusters, this relative advantage of Bayesian RLCM persists under data replications.

**Figure 3:**
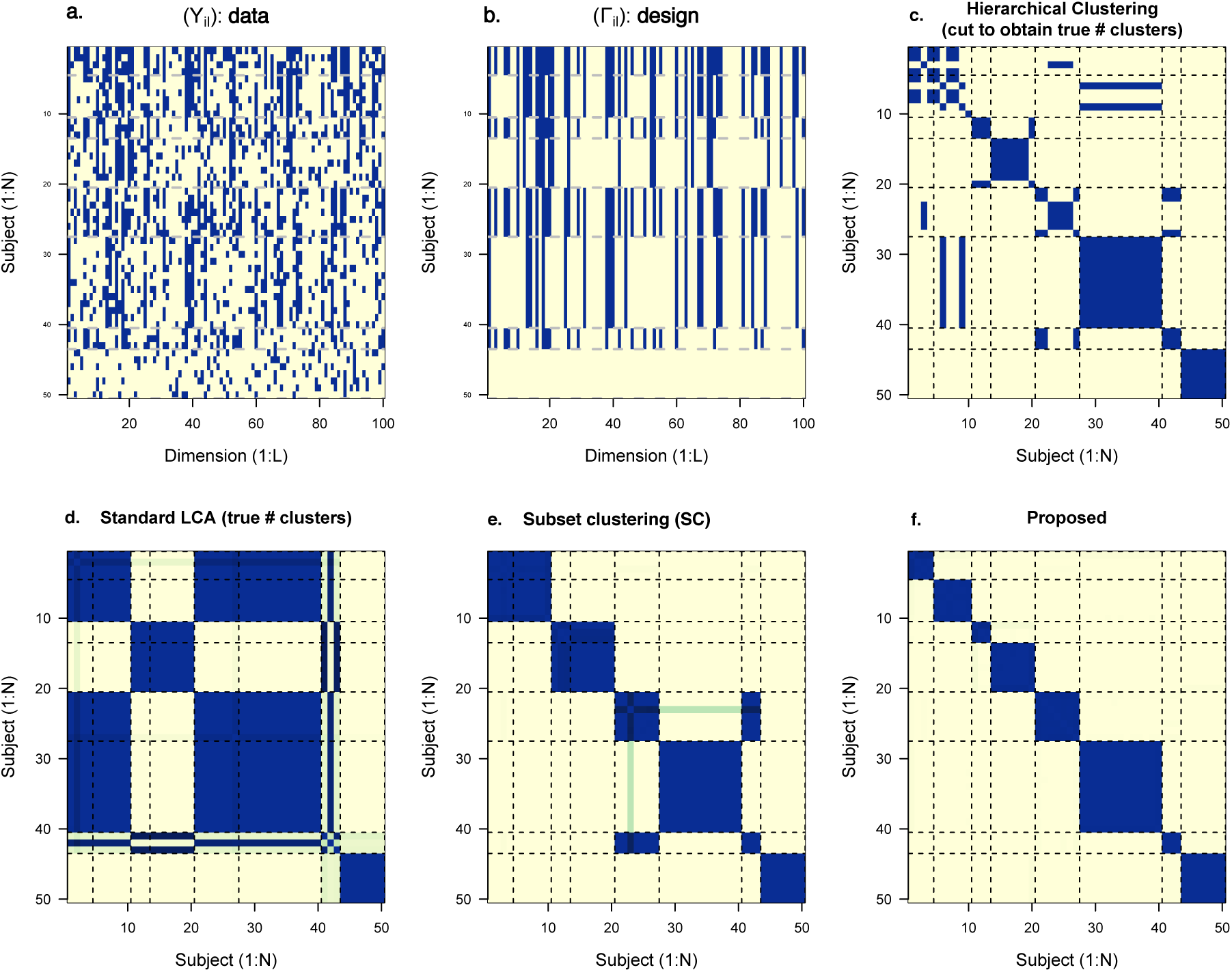
In the 100-dimension multivariate binary data example (a), the eight classes differ with respect to subsets of measured features (b). In (c) HC, we indicate coclustering by filled cells. The true clusters are separated (dashed grids) and ordered according to the truth; (d, e, f): For Bayesian LCA, RLCM and subset clustering (SC), we plot the posterior co-clustering probability matrix 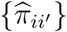 for *N* observations. Filled blocks on the main diagonal only indicate perfect recovery of the true clusters; Blank cells within the main diagonal blocks indicate true cluster being split and blue cells in the off-diagonal blocks indicate two observations being incorrectly co-clustered. Bayesian RLCM accounts for measurement errors, selects the relevant feature subsets and filters the subsets by a low-dimensional model (1) and therefore yields superior clustering results. The clustering results are based on a randomly generated data set for illustration. Cluster recovery by RLCM is not always perfect. The competing methods may show different relative performances under model mis-specifications (see Section 4.1). (This figure appears in color in the electronic version of this article, and any mention of color refers to that version.)

#### Simulation 2: Assess clustering performance under various parameter settings with replications

The ability of Bayesian RLCM in recovering the true clusters varies by the sparsity level (*s*) in each machine, level of measurement errors 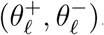, mixing weights and sample sizes (*N*) (the leftmost boxes in groups of four in Figure 4). Firstly, clustering performance improves by increasing the sparsity level in each machine from *s* = 10% to 20% (compare the 1st and 3rd, 2nd and 4th RLCM boxplots in each panel of Figure 4). In the context of our motivating example, given a fixed number of protein landmarks *L*, patients will be more accurately clustered if each machine comprises more component proteins. This observation is also consistent with simulation studies conducted in the special case of *Q* = *I*_*L*_ (Hoff, 2005, *Table 1). For a given s*, a larger *L* means a larger number of relevant features per machine and leads to better cluster recovery. Increasing *L* from 50 to 400, the mean aRI (averaged over replications) increases from 0.7 to 0.98 at the sparsity level *s* = 10%, 0.88 to 0.99 under *s* = 20%. Secondly, more accurate clustering results under larger discrepancies between 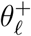 and 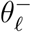. The aRI averaged over replications is higher under 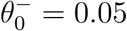 than 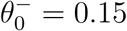 over all combinations of other parameters. Finally we observe mixed relative performances at distinct sample sizes as a result of two competing factors as the sample size increases: more precise estimation of measurement error parameters that improve clustering and a larger space of clusterings. Additional comparisons are in Figure S1 of the Supporting Information.

**Figure 4:**
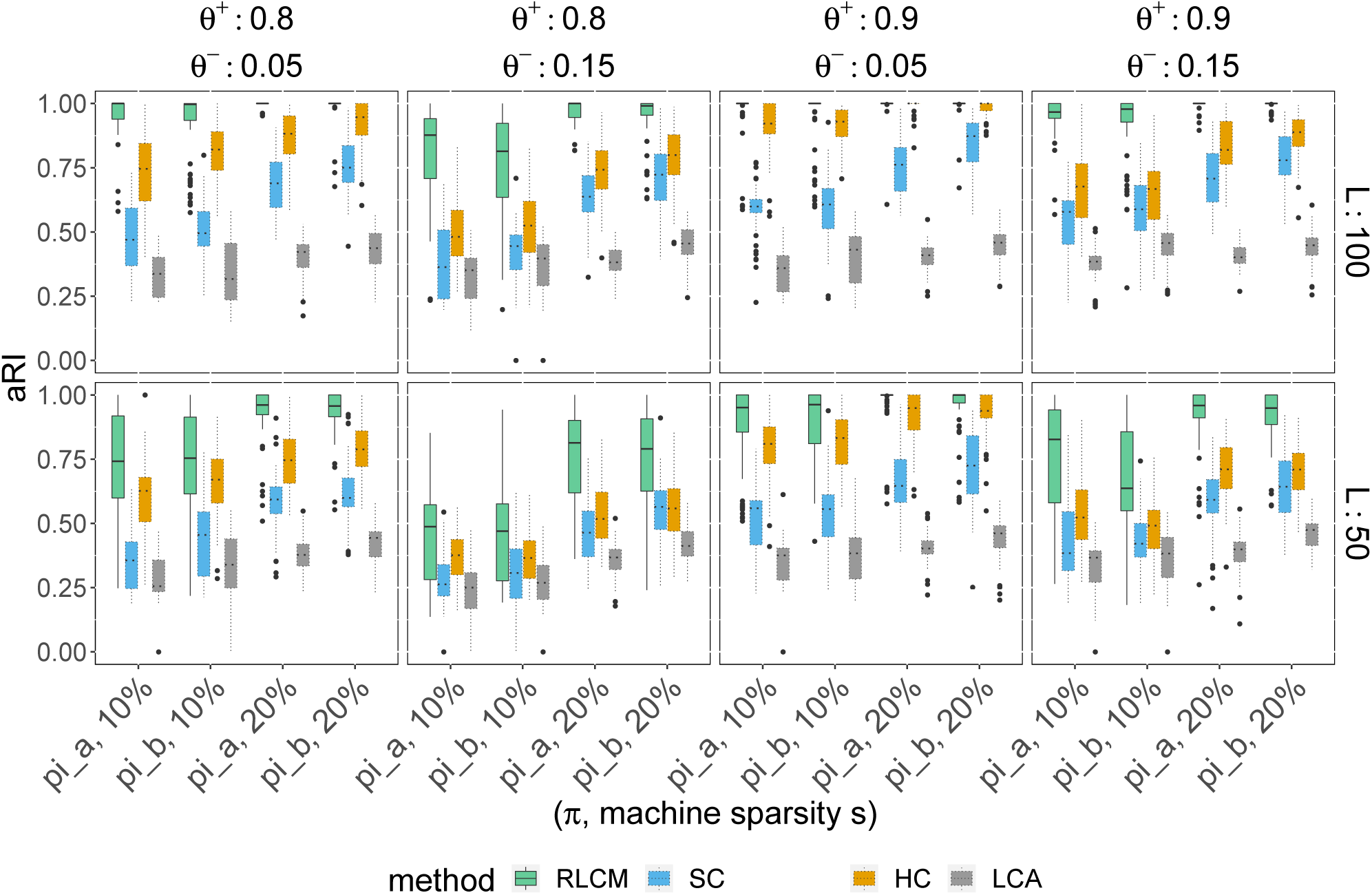
Based on *R* = 60 replications for each parameter setting, from the left to the right in each group of four boxplots, Bayesian RLCM (boxplots with solid lines) most accurately recovers the true clusters compared to subset clustering (SC) hierarchical clustering (HC) and traditional Bayesian latent class analysis (LCA). (This figure appears in color in the electronic version of this article, and any mention of color refers to that version.)

Compared to three common alternatives, the Bayesian RLCM on average most accurately recovers the clusters. Bayesian RLCM produces the highest aRIs (boxes with solid lines) compared to others (boxes with dotted lines). For example, under 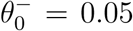, the ratio of the mean aRIs (averaged over replications) for Bayesian RLCM relative to subset clustering is 2.06, 2.04, 1.88, 1.71 for the sample-size-to-dimension ratios *N/L* = 1, 0.5, 0.25, 0.125, respectively. As another example, under a higher 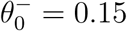, the relative advantage of Bayesian RLCM to HC narrows as shown by the smaller aRI ratios 1.23, 1.62, 1.49, 1.16. Web Appendix A5.3 provides further discussion about the three alternative methods.

#### Simulation 3

Competing methods are evaluated under small and large degrees of model misspecifications for fairer comparisons. We consider two simple representative scenarios: 1) randomly perturbing only the final layer of the model to have more than two levels of response probabilities; 2) removing any restriction on how the clusters are connected to the observables by assuming general LCMs. See Web Appendix A5.1 for specific parameter values. We observe that in the first set with relatively minor misspecifications, the proposed model is still competitive (Figure S2 in the Supporting Information). The proposed approach can be viewed as regularization by shrinking class-specific response probabilities towards the assumed constraint. RLCM expectedly performs less well under larger magnitudes of random perturbations. In the second set, the main model is not as flexible as a general LCM resulting in less competitive clustering performance (Figure S3 in the Supporting Information). Large degrees of model misspecifications may hurt clustering performance. Our practical suggestion is to perform careful model checking which we illustrate in Section 4.2.

### 4.2 Protein Data Application for Estimating Scleroderma Patient Subsets

The applied goal is to estimate autoimmune disease patient clusters via reconstructing machine components. We seek to identify components of the machines and to quantify the variations in their occurrence among individuals and estimate patient subsets. The binary responses ***Y***_*i*_ indicate the observed presence or absence of proteins at equi-spaced molecular weight landmarks as produced by a preprocessing method (Wu et al., 2019) applied to GEA data. We ran 4 gels, each loaded with immunoprecipitations (IPs) performed using sera from 19 different patients, and one reference lane. All sera were from scleroderma patients with cancer, and were all negative for the three most common autoantibodies found in scleroderma (anti-RNA polymerase III, anti-topoisomerase I, and anti-centromere). The IPs were loaded in random order on each gel; the reference sample is comprised of known molecules of defined sizes (molecular weights) and was always loaded in the first lane. The left panel in Figure 5 shows for each sample lane (labeled in the left margin; excluding the reference lanes) the binary responses indicating the observed presence or absence of autoantibodies at *L* = 50 landmarks.

**Figure 5:**
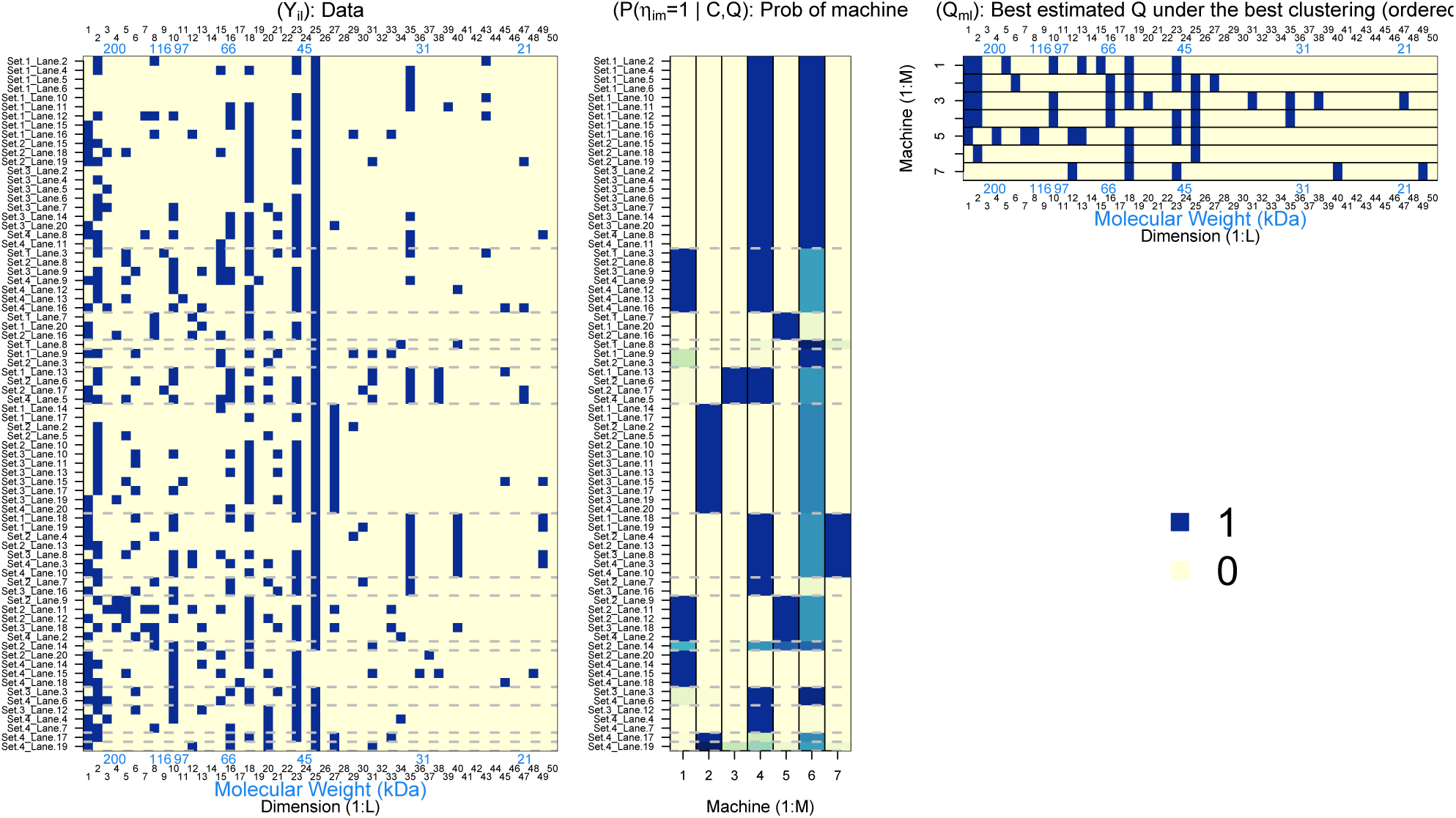
Results for GEA data. *Left* : Aligned data matrix for band presence or absence; row for 76 serum lanes, reordered into optimal estimated clusters (not merged) 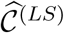 separated by gray horizontal lines “—–”; columns for *L* = 50 protein landmarks. A blue vertical line “|” indicates a band; *Middle*: lane-machine matrix for the probability of a lane (serum sample) having a particular machine. The blue cells correspond to high probability of having a machine in that column. Smaller probabilities are shown in lighter blue;. *Right* : The estimated machine profiles. Here seven estimated machines are shown, each with component proteins shown by a blue bar “| “. (This figure appears in color in the electronic version of this article, and any mention of color refers to that version.)

Patients differ in protein presence or absence patterns at the protein landmarks. Eleven out of *L* = 50 aligned landmarks are absent among the patients tested. The rest of the landmarks are observed with prevalences between 1.3% and 94.7%. The GEA technologies are known to be highly specific and sensitive for nearly all proteins studied in this assay so we specify the hyperparameters in the Beta priors by *N*_1_ = 10, *a*_1_ = 0.9, *N*_0_ = 100, *a*_0_ = 0.01 and conducted sensitivity analyses varying these hyperparameter values.

In this application, the scientists had previously identified and independently verified through additional protein chemistry the importance of a small subset of protein bands in determining clusters among a subset of subjects. They proposed that these subjects should be grouped together. We therefore fitted the Bayesian RLCM without further splitting these partial clusters *𝒞*^(0)^ so that the number of scientific clusters visited by the MCMC chain has an upper bound 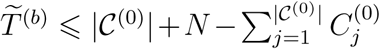, where 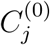 counts the number of observations in the initial cluster *j*. We fitted models and compared the results under multiple working truncation levels *M* ^*†*^ = 8, 9, …, 15 and obtained identical clustering results.

Figure 5 shows the observations grouped by the RLCM-estimated clusters (not merged) 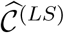 (left), the estimated *Q*-matrix 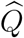 (right), and the conditional posterior probabilities of the machines 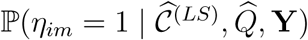 (middle).

The information for estimating matrix *Q* comes from the observed marginal associations (positive or negative) among the protein landmarks. Landmark protein pairs observed with positive association tend to be placed in the same estimated machine. For example, Landmarks 4, 7 and 8 appear together in Machine 5. Subjects either have all three landmarks or none at all, which induces strong positive pairwise associations among these landmarks. Indeed, the estimated log odds ratio (LOR) is 3.13 (standard error 1.16) for Landmark 4 versus 7, 2.21 (s.e., 0.98) for Landmark 4 versus 8, and 2.92 (s.e. 1.2) for Landmark 7 versus 8. The observed negative marginal associations between two landmarks suggest existence of machines with discordant landmarks. For example, Landmarks 10 and 27 are rarely estimated to be present or absent together in a subject as a result of 1) estimated machines with discordant landmarks and 2) subject-specific machine assignments. First, the model estimated that Landmark 10 (in Machine Set A: 1, 3 and 4) belongs to machines not having Landmark 27 (it is in Machine Set B: 2). Second, with high posterior probabilities, most observations have machines from one of, not both Set A and B hence creating discordance (high posterior probability ℙ (Γ_*i*10_ ≠ Γ_*i*27_ | **Y**)). In the presence of measurement errors, strong negative marginal association results (observed LOR for Landmark 10 versus 27: −1.98, s.e. 0.8).

Our algorithm also directly infers the number of scientific clusters in the data given an initial partial clustering *𝒞*^(0)^. The marginal posterior of the number of scientific clusters 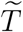 can be approximated by empirical samples of 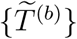 which result in a posterior median of 12 (95% credible interval: (8, 16); Figure S4 in the Supporting Information). The advantage of Bayesian RLCM is the posterior inference about both the clusters and the distinct latent state variables ***η***_*i*_ interpreted based on the inferred *Q* matrix. The middle panel of Figure 5 shows that clusters differ in their posterior probabilities of having each of the estimated machines. Among 76 subjects analyzed, 23 of them have greater than 95% posterior probabilities of having both Machine 4 and 6. Finally, a group of seven subjects are estimated to be enriched with Machine 4 and 7. This is expected because the two machines have one or more of the Landmarks 35, 40 and 49 (33, 27 and 18 kDa bands, respectively) and together explain the distinctive observed combination of the raw bands in the seven subjects. Such inference about ***η***_*i*_ is not available based on hierarchical clustering or traditional latent class analysis.

We performed posterior predictive checking to assess model fit (Gelman et al., 1996). At each MCMC iteration, given the posterior sample of model parameters, we simulated a data set of the same size as the original set. For each replicated data set, we compute the marginal means and marginal pairwise log odds ratios (0.5 adjustment for zero counts). Across all replications, we compute the 95% posterior predictive confidence intervals (PPCI) defined by the 2.5% and 97.5% quantiles of the PPD. All the observed marginal means are covered by their respective PPCIs; The 95% PPCIs cover all but 24 of 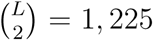 landmark pairs of observed pairwise log odds ratios (see Figure S5 and S6 in the Supporting Information). The proposed model adequately fits the GEA data.

## 5. Discussion

Modern scientific technologies give rise to measurements of varying precision and accuracy that are better targeted at the underlying state variables than ever before. In this paper we have focused on finite-sample Bayesian inference of an RLCM for analyzing dependent binary data. The primary advantage of such models lies in their expressive characterization of the between-class differential errors structured to respect specific scientific context about the biological and measurement processes. Using simulations and real data analysis, we studied the clustering of observations with an unknown number of clusters, uncertainty assessment of the clustering and the prediction of individual latent states. We develop and apply a MCMC algorithm for Bayesian RLCMs. The proposed method addresses inferential issues unique to mixture models with discrete component parameters and jointly infers the number of clusters, the design matrix Γ and other model parameters. We have illustrated its advantage through simulations relative to three commonly used binary-data clustering methods. Finally, viewed from a regularization perspective, the inferential procedure automatically selects subsets of features for each latent class and filters them through a low-dimensional model that shrinks class-specific response probability estimates toward one that represents the scientific structure and improves our ability to accurately estimate clusters.

RLCMs decompose the variation among multivariate binary responses into structure that reflects constraints in particular scientific context and stochastic variation without a known explanation. In our motivating example, it is certainly likely that there is some variability related to measurement errors. However, it is also highly likely that there are systematic biological and biochemical processes not included in the structural part because they are unknown to us today. RLCM analyses can be a useful tool in the effort to uncover the unknown structure. One approach would be to show that the latent classes are diagnostic of specific diseases. Another is that we might uncover a novel mechanism by defining distinct patterns of the same autoantigen machine in patients with the same disease or potentially in patients with different diseases that target the same machines. Though the present paper focused on an example in medicine, the developed method and algorithms apply to many problems in psychology and epidemiology.

We are currently studying a few potentially-useful extensions. First, nested partially LCMs (Wu et al., 2017) incorporate local dependence and multiple sensitivity parameters that would improve the utility of Bayesian RLCMs. Second, because the algorithm involves iterating over subjects to find clusters, the computational time increases with the number of subjects *N*. Divide-Cluster-Combine schemes that estimate clusters in subsamples which are then combined may improve the computational speed at the expense of the approximation introduced by the multi-stage clustering (Ni et al., 2020). Third, in applications where the clustering of multivariate binary data comprises an important component of a hierarchical Bayesian model with multiple components, the posterior uncertainty in clustering propagates into other parts of the model and can be integrated into posterior inference of other model parameters (e.g., Jacob et al., 2017). Finally, under significant deviations from model assumptions, the constraints in Γ may lead to less competitive clustering performance. In real data applications, scientific context for clustering and careful model checking are therefore critical. The degree to which the actual data generating mechanism deviates from the assumed model can be characterized by general RLCM formulations such as the all-effect model (Gu and Xu, 2019a, Equation (5)). Model selection under such settings may help with clustering performance. We leave this for future work.

## Supporting information

The Supplementary Materials contain the technical details, extra simulation results and figures referenced in Main Paper.

## Acknowledgment

The research is supported in part by a gift from the Jerome L. Greene Foundation and by the Patient-Centered Outcomes Research Institute (PCORI) Award (ME-1408-20318), National Institutes of Health (NIH) grants R01AR073208, P30AR070254 and P30CA046592. We are grateful for comments from the Editor, Associate Editor and three reviewers that greatly improved the presentation. We also thank Gongjun Xu, Peter Hoff and Jian Kang for helpful discussions.

## Supporting Information

Web Appendices, Tables, and Figures referenced in Sections 2.3, 3.1 and 4 are available with this paper at the *Biometrics* website on Wiley Online Library. R package “rewind” is freely available at https://github.com/zhenkewu/rewind. Another R package “spotsgear” (https://github.com/zhenkewu/spotgear) is used for data preprocessing in Section 4.2.

## Notes

### Competing Interest Statement

The authors have declared no competing interest.

### Summary of Updates

Main text is now updated.

## References

Albert, J. H. and Chib, S. (1993). Bayesian analysis of binary and polychotomous response data. Journal of the American statistical Association 88, 669–679.

Chiu, C.-Y., Douglas, J. A., and Li, X. (2009). Cluster analysis for cognitive diagnosis: theory and applications. Psychometrika 74, 633–665.

Dahl, D. B. (2006). Model-based clustering for expression data via a Dirichlet process mixture model. In Do, K. A., Müller, P., and Vannucci, M., editors, Bayesian Inference for Gene Expression and Proteomics, pages 201–218. Cambridge University Press, New York.

Dunson, D. and Xing, C. (2009). Nonparametric Bayes modeling of multivariate categorical data. Journal of the American Statistical Association 104, 1042–1051.

Garrett, E. and Zeger, S. (2000). Latent class model diagnosis. Biometrics 56, 1055–1067.

Gelfand, A. E. and Smith, A. F. (1990). Sampling-based approaches to calculating marginal densities. Journal of the American Statistical Association 85, 398–409.

Gelman, A., Meng, X.-L., and Stern, H. (1996). Posterior predictive assessment of model fitness via realized discrepancies. Statistica Sinica 6, 733–760.

Ghahramani, Z. and Griffiths, T. L. (2006). Infinite latent feature models and the Indian buffet process. In Advances in Neural Information Processing Systems, pages 475–482.

Goodman, L. (1974). Exploratory latent structure analysis using both identifiable and unidentifiable models. Biometrika 61, 215–231.

Green, P. J. (1995). Reversible jump Markov chain Monte Carlo computation and Bayesian model determination. Biometrika 82, 711–732.

Gu, Y. and Xu, G. (2019a). Learning attribute patterns in high-dimensional structured latent attribute models. Journal of Machine Learning Research 20, 1–58.

Gu, Y. and Xu, G. (2019b). The sufficient and necessary condition for the identifiability and estimability of the DINA model. Psychometrika 84, 468–483.

Hoff, P. D. (2005). Subset clustering of binary sequences, with an application to genomic abnormality data. Biometrics 61, 1027–1036.

Hubert, L. and Arabie, P. (1985). Comparing partitions. Journal of Classification 2, 193–218.

Jacob, P. E., Murray, L. M., Holmes, C. C., and Robert, C. P. (2017). Better together? statistical learning in models made of modules. arXiv preprint 1708.08719.

Jain, S. and Neal, R. M. (2004). A split-merge Markov chain Monte Carlo procedure for the Dirichlet process mixture model. Journal of Computational and Graphical Statistics 13, 158–182.

Junker, B. W. and Sijtsma, K. (2001). Cognitive assessment models with few assumptions, and connections with nonparametric item response theory. Applied Psychological Measurement 25, 258–272.

Kadane, J. (1975). The role of identification in Bayesian theory. In Fienberg, S. and Zellner, A., editors, Studies in Bayesian Econometrics and Statistics, chapter 5.2, pages 175–191. North-Holland, Amsterdam.

Lazarsfeld, P. F. (1950). The logical and mathematical foundations of latent structure analysis. In Stouffer, S., editor, The American Soldier: Studies in Social Psychology in World War II, volume IV, pages 362–412. Princeton University Press, Princeton, NJ.

Lee, D. D. and Seung, H. S. (1999). Learning the parts of objects by non-negative matrix factorization. Nature 401, 788–791.

Liu, J. S. (1996). Peskun’s theorem and a modified discrete-state Gibbs sampler. Biometrika 83, 681–682.

McCullagh, P., Yang, J., et al. (2008). How many clusters? Bayesian Analysis 3, 101–120.

Meeds, E., Ghahramani, Z., Neal, R. M., and Roweis, S. T. (2007). Modeling dyadic data with binary latent factors. In Advances in Neural Information Processing Systems, pages 977–984.

Miettinen, P., Mielikäinen, T., Gionis, A., Das, G., and Mannila, H. (2008). The discrete basis problem. IEEE Transactions on Knowledge and Data Engineering 20, 1348–1362.

Miller, J. W. and Harrison, M. T. (2018). Mixture models with a prior on the number of components. Journal of the American Statistical Association 113, 340–356.

Ni, Y., Müller, P., Diesendruck, M., Williamson, S., Zhu, Y., and Ji, Y. (2020). Scalable Bayesian nonparametric clustering and classification. Journal of Computational and Graphical Statistics 29, 53–65.

Ni, Y., Müller, P., and Ji, Y. (2019). Bayesian double feature allocation for phenotyping with electronic health records. Journal of the American Statistical Association, To Appear.

Nobile, A. and Fearnside, A. T. (2007). Bayesian finite mixtures with an unknown number of components: The allocation sampler. Statistics and Computing 17, 147–162.

Rosen, A. and Casciola-Rosen, L. (2016). Autoantigens as partners in initiation and propagation of autoimmune rheumatic diseases. Annual Review of Immunology 34, 395–420.

Rukat, T., Holmes, C. C., Titsias, M. K., and Yau, C. (2017). Bayesian boolean matrix factorisation. In International Conference on Machine Learning, pages 2969–2978.

Teh, Y. W., Grür, D., and Ghahramani, Z. (2007). Stick-breaking construction for the Indian buffet process. In Artificial Intelligence and Statistics, pages 556–563.

Templin, J. L. and Henson, R. A. (2006). Measurement of psychological disorders using cognitive diagnosis models. Psychological Methods 11, 287–305.

Vermunt, J. K. and Magidson, J. (2002). Latent class cluster analysis. Applied Latent Class Analysis 11, 89–106.

Wu, Z., Casciola-Rosen, L., Shah, A., Rosen, A., and Zeger, S. L. (2019). Estimating autoantibody signatures to detect autoimmune disease patient subsets. Biostatistics 20, 30–47.

Wu, Z., Deloria-Knoll, M., Hammitt, L. L., and Zeger, S. L. (2016). Partially latent class models for case–control studies of childhood pneumonia aetiology. Journal of the Royal Statistical Society: Series C (Applied Statistics) 65, 97–114.

Wu, Z., Deloria-Knoll, M., and Zeger, S. L. (2017). Nested partially latent class models for dependent binary data; estimating disease etiology. Biostatistics 18, 200–213.

Xu, G. (2017). Identifiability of restricted latent class models with binary responses. The Annals of Statistics 45, 675–707.

Xu, G. and Shang, Z. (2018). Identifying latent structures in restricted latent class models. Journal of the American Statistical Association 113, 1284–1295.

Zhang, Z., Li, T., Ding, C., and Zhang, X. (2007). Binary matrix factorization with applications. In Seventh IEEE International Conference on Data Mining (ICDM 2007), pages 391–400.

