## Supplementary material for "A Bayesian Approach to Restricted Latent Class Models for Scientifically-Structured Clustering of Multivariate Binary Outcomes": The Supplementary Materials contain the technical details, extra simulation results and figures referenced in Main Paper.


#### A1 Prior Distribution (8)-(10) in the Main Paper

##### A1.1 The Role of Hyperparameters $c_1$ and $c_2$

By Beta-Bernoulli conjugacy, we integrate the joint distribution in (8)-(9) in Main Paper $[\{\boldsymbol{\alpha}_k, k = 1, \ldots, K\} \mid \boldsymbol{p}][\boldsymbol{p} \mid c_1, c_2]$ over $\boldsymbol{p}$ to obtain the marginal prior:

$$pr(\{\boldsymbol{\alpha}_k, k = 1, \ldots, K\} \mid c_1, c_2) = \prod_{m=1}^{M} \frac{(c_1c_2/M)\Gamma(n_{m1} + c_1c_2/M)\Gamma(K - n_{m1} + c_2)}{\Gamma(K + c_2 + c_1/M)}, \quad \text{(S1)}$$

where $\Gamma(\cdot)$ is the Gamma function and $n_{m1} = \sum_{k=1}^{K} \alpha_{km}$, $m = 1, \ldots, M$. Holding $c_2$ constant, the prior average number of positives among $\boldsymbol{\alpha}_k$ decreases with $c_1$. Holding $c_1$ constant, the latent state vectors, $\boldsymbol{\alpha}_k$ and $\boldsymbol{\alpha}_{k'}$, $k \neq k'$, become *a priori* increasingly similar as $c_2$ decreases. In fact, the prior probability of $\mathbb{P}[\alpha_{km} = \alpha_{k'm} \mid k \neq k', c_1, c_2] = \mathbb{E}\{p_m^2 + (1 - p_m)^2 \mid c_1, c_2\} = 1 - 2\frac{c_1}{c_1+M}\left(1 - \frac{c_1c_2+M}{c_1c_2+c_2M+M}\right)$ approaches one when $c_2$ approaches zero.

#### A1.2 On Merging Clusters with Identical Draws of States

Define "scientific clusters" $\widetilde{\mathcal{C}}$ by merging clusters associated with identical latent states. That is,

$$\widetilde{\mathcal{C}} = \left\{\{i : \boldsymbol{\eta}_i = \widetilde{\boldsymbol{\alpha}}_k, k = 1, \ldots, \widetilde{T}\right\}$$

where $\{\widetilde{\boldsymbol{\alpha}}_k, k = 1, \ldots, \widetilde{T}\}$ collects $\widetilde{T}(\leq T)$ unique patterns among $\{\boldsymbol{\alpha}_k, k = 1, \ldots, T\}$ that are present in the sample. Let $\mathcal{M} : \{\boldsymbol{\eta}_i = \boldsymbol{\alpha}_{Z_i}, i = 1, \ldots, N\} \mapsto \widetilde{\mathcal{C}}$ represent this merge operation, i.e., $\widetilde{\mathcal{C}} = \mathcal{M}(H)$, where $H = \{\boldsymbol{\eta}_i, i = 1, \ldots, N\}$.

Define partial ordering " $\preceq$ " over partitions $\mathcal{C}_1 \preceq \mathcal{C}_2$ if for any $C_1 \in \mathcal{C}_1$, one can find a $C_2 \in \mathcal{C}_2$ satisfying $C_1 \subseteq C_2$. We have $\mathcal{C} \preceq \widetilde{\mathcal{C}}$, i.e., $\widetilde{\mathcal{C}}$ is coarser than $\mathcal{C}$. Our procedure for obtaining clusters $\widetilde{\mathcal{C}}$ differs from the kind of mixture models that distinct $Z_i$ values with probability one correspond to distinct component parameters sampled from a continuous base measure (e.g., Miller and Harrison, 2017, Proof of Theorem 4.2).

We specified priors on $\mathcal{C}$. And for each cluster, we have $\{\boldsymbol{\alpha}_k, k = 1, \ldots, T\}$, where $T \leq K$ and excludes empty component(s). We can then merge $\mathcal{C}$ to obtain $\widetilde{\mathcal{C}}$. Therefore a prior on $\mathcal{C}$ induces a prior on $\widetilde{\mathcal{C}}$. Setting $c_2 = 1$, we have

$$p(\widetilde{\mathcal{C}} \mid c_1, \gamma) = \sum_{\mathcal{C}:\mathcal{C}\preceq\widetilde{\mathcal{C}}} p(\widetilde{\mathcal{C}} \mid \mathcal{C}, c_1) \cdot p(\mathcal{C} \mid \gamma) \tag{S2}$$

$$= \sum_{\mathcal{C}:\mathcal{C}\preceq\widetilde{\mathcal{C}}} \binom{2^M}{\widetilde{T}} (\widetilde{T})! \left\{ \int p(\{\boldsymbol{\alpha}_k, k = 1, \ldots, T\} \mid \boldsymbol{\mathcal{S}}, \boldsymbol{p}) p(\boldsymbol{p} \mid c_1) \mathrm{d}\boldsymbol{p} \right\} \cdot p(\mathcal{S} \mid \gamma) \cdot T!, \tag{S3}$$

where $\mathcal{S} = \{S_1, \ldots, S_T\}$ is a ordered partition of $N$ subjects, obtained by randomly ordering parts or blocks of $\mathcal{C}$ uniformly over $T!$ possible choices and $p(\mathcal{S} \mid \gamma) \cdot T! = p(\mathcal{C} \mid \gamma)$.

### A2 Marginal Likelihood $g(C)$

Given assignment of subjects to blocks in $\mathcal{C}$, the model likelihood in a cluster $C \in \mathcal{C}$ is

$$pr\left(\mathbf{Y}_C \mid \boldsymbol{\alpha}, \boldsymbol{\theta}^+, \boldsymbol{\theta}^-, Q\right) = \prod_{\ell:\Gamma(Q,\boldsymbol{\alpha})_\ell=0} \{\theta_\ell^-\}^{n_{k\ell1}} \left(1 - \theta_\ell^-\right)^{n_{k\ell0}} \cdot \prod_{\ell:\Gamma(Q,\boldsymbol{\alpha})_\ell=1} \{\theta_\ell^+\}^{n_{k\ell1}} \left(1 - \theta_\ell^+\right)^{n_{k\ell0}}, \tag{S4}$$

where $n_{k\ell1} = \sum_{i\in C} Y_{i\ell}$ and $n_{k\ell0} = \sum_{i\in C}(1 - Y_{i\ell})$ are the number of positive and negative responses at dimension $\ell$ for subjects in cluster $C$. We obtain the marginal likelihood $g(C)$ by integrating out latent states $\boldsymbol{\alpha}$ in (S4):

$$g(C) = \sum_{\boldsymbol{\alpha}\in\{0,1\}^M} pr\left(\mathbf{Y}_C \mid \boldsymbol{\alpha}, \boldsymbol{\theta}^+, \boldsymbol{\theta}^-, Q\right) \mathbb{P}(\boldsymbol{\alpha}_k = \boldsymbol{\alpha} \mid \boldsymbol{p}), \tag{S5}$$

where $\mathbb{P}(\boldsymbol{\alpha}_k = \boldsymbol{\alpha} \mid \boldsymbol{p}) = \prod_{m=1}^M p_m^{\alpha_m}(1 - p_m)^{1-\alpha_m}$.

#### A3 Split-Merge Update

We adapt an existing recipe designed for models with priors conjugate to the component-specific parameters (Jain and Neal, 2004). The goal of split-merge updates is to make global changes to cluster configuration followed by further refinement of clusters via Gibbs update one subject at a time. We sketch a complete round of split-merge update in the following; see Jain and Neal (2004) for full details.

Step i: Randomly choose two observations $i$ and $j$. Let $S = \{i' : Z_{i'} = i \text{ or } j, i' = 1, \ldots, N\}$.

Step ii: Based on (12) in the Main Paper, assign subject $k$ in $S \setminus \{i, j\}$ to either the cluster of $i$ or $j$ with probability $\mathbb{P}(Z_k = z \mid \text{others})$:

$$\frac{(|C_z| + \gamma) g(C_z \cup \{k\})/g(C_z)}{(|C_{Z_i}| + \gamma) g(C_{Z_i} \cup \{k\})/g(C_{Z_i}) + (|C_{Z_j}| + \gamma) g(C_{Z_j} \cup \{k\})/g(C_{Z_j})}, \tag{S6}$$

for $z \in \{Z_i, Z_j\}$. Repeat the intermediate Gibbs scan for $r = 5$ times and obtain $\boldsymbol{Z}^{\text{launch}}$.

Step iii: Perform a final Gibbs scan restricted to observations $S \setminus \{i, j\}$ using (S6), resulting in updated clusters as the proposal states to be used in a Metroplis-Hasting step which we denote by $\boldsymbol{Z}^{\text{cand}}$. Compute the proposal densities $q(\boldsymbol{Z}^{\text{cand}} \mid \boldsymbol{Z})$ and $q(\boldsymbol{Z} \mid \boldsymbol{Z}^{\text{cand}})$. The details are in Jain and Neal (2004). For the non-trivial cases, the proposal densities depend on the random launch state $\boldsymbol{Z}^{\text{launch}}$ and are products of Gibbs update densities in (S6). $\boldsymbol{Z}^{\text{launch}}$ appears in the proposal densities, because it indexes the transition kernel to $\boldsymbol{Z}^{\text{cand}}$.

Step iv: Accept or reject the proposed clustering $\boldsymbol{Z}^{\text{cand}}$ with acceptance probability computed from prior ratio (based on two sets of clusters induced by $\boldsymbol{Z}^{\text{cand}}$ vs $\boldsymbol{Z}^{\text{launch}}$), likelihood ratio (given clusters $\boldsymbol{Z}^{\text{cand}}$ vs $\boldsymbol{Z}^{\text{launch}}$ and other population parameters), ratio of proposal densities (from Step iii). See Jain and Neal (2004) for the general recipe of computing the acceptance probability.

Step v: Perform one complete Gibbs scan (12) in the Main Paper for all observations to refine the current state of cluster indicators.

The above is referred to as $(5, 1, 1)$ split-merge update where 5 intermediate Gibbs scans are used to reach launch states $\boldsymbol{Z}^{\text{launch}}$, one Metroplis-Hasting step to accept or reject a candidate clustering $\boldsymbol{Z}^{\text{cand}}$, and one final complete Gibbs scan for all observations to refine the newly obtained cluster (Jain and Neal, 2004).

#### A4 Convergence Check

In simulations and data analysis, we ran three MCMC chains each with a burn-in period of $10,000$ iterations followed by $10,000$ iterations stored for posterior inference. We look for potential non-convergence in terms of Gelman-Rubin statistic (Brooks and Gelman, 1998) that compares between-chain and within-chain variances for each model parameter where a large difference ($R_c > 1.1$) indicates non-convergence; We also used Geweke's diagnostic (Geweke and Zhou, 1996) that compare the observed mean for each unknown variable using the first 10% and the last 50% of the stored samples where a large $Z$-score indicates non-convergence ($|Z| > 2$). In our simulations and data analyses, we observed fast convergence (many satisfied convergence criteria within $2,000$ iterations) that led to well recovered clusters and $Q$ matrices (results not shown here).

### A5 Details about Simulation Studies

#### A5.1 Simulation Setup

*Simulation 1.* We set $N = 50$, $L = 100$ and $M = 3$. We randomly generate a matrix $Q$ ($M$ by $L$) where each row has on average $s = 20\%$ non-zero elements: $Q_{m\ell} \stackrel{i.i.d}{\sim} \mathsf{Bernoulli}(0.2), \ell = 1, \ldots, L$. In the rare event where a random $Q \notin \mathcal{Q}$ defined by (4) in the Main Paper, we randomly permute pairs of elements in $Q_{m*}$ until $Q \in \mathcal{Q}$. We draw latent states for each observation independently according to $\boldsymbol{\eta}_i \stackrel{d}{\sim} \mathsf{Categorical}\left(\mathcal{A}; \boldsymbol{\pi}_0 = \boldsymbol{\pi}_b\right)$ where

$$\boldsymbol{\pi}_0 = \{\mathbb{P}(\boldsymbol{\eta}_i = (0,0,0), (1,0,0), (0,1,0), (1,1,0), (0,0,1), (1,0,1), (0,1,1), (1,1,1))\},$$

and

$$\boldsymbol{\pi}_b = (1/6, 1/6, 1/6, 1/6, 1/12, 1/12, 1/12, 1/12).$$

We assume the response probabilities shift between two levels $\theta_\ell^+ = 0.8$ and $\theta_\ell^- = 0.15$. The distinct subsets of features where shifts occur define eight classes $|\mathcal{A}| = 8 = (2^M)$, which upon enumeration by observation gives an $N$ by $L$ design matrix $\Gamma$.

*Simulation 2.* We simulated $R = 60$ replication data sets for each of $1,920$ combinations of (#features, sample size, sensitivity, 1-specificity, mixing weights, sparsity level): $(L, N, \theta_0^+, \theta_0^-, \boldsymbol{\pi}_0, s) \in \{50, 100, 200, 400\} \otimes \{50, 100, 200\} \otimes \{0.8, 0.9\} \otimes \{0.05, 0.15\} \otimes \{\boldsymbol{\pi}_a = (\frac{1}{8}, \ldots, \frac{1}{8}), \boldsymbol{\pi}_b = (\frac{1}{6}, \ldots, \frac{1}{6}, \frac{1}{12}, \ldots, \frac{1}{12})\} \otimes \{10\%, 20\%\}$. The parameter values are designed to mimic what would be expected in the motivating example.

*Simulation 3.* We further compare clustering performance of the competing methods under two simple representative sets of data generating mechanisms (DGM) with small and large degrees of departures from model (1)-(3) in the main paper. The choices are guided by their scientific relevance to the scleroderma application. The first set only perturbs the final layer of the model. It assumes the same scientific structure as the main model and the same parameter values in Simulation 2 except the measurement error parameters. This is to mimic hypothetical variation in the GEA experiment conditions which might be missed by the assumed model. More specifically, for subject $i$ in class $k$, we set $\theta_{i\ell} \sim \theta_\ell^+ + U_{k\ell}$ if $\Gamma_{i\ell} = 1$ and $\theta_{i\ell} \sim \theta_\ell^- + V_{k\ell}$ if $\Gamma_{i\ell} = 0$, where $U_{k\ell} \sim \mathsf{Uniform}(-0.095, 0.095)$ and $V_{k\ell} \sim \mathsf{Uniform}(-0.045, 0.045)$ independently for class $k = 1 \ldots, K$ and feature $\ell = 1, \ldots, L$. We set other parameters prior to perturbation the same as in Simulation 2. The second set is the classical LCM. Different from any RLCM, it does not specify any structure on how the clusters are linked to the observables so the classes have flexible response probability profiles. We simulated data under three- and six-class LCMs to investigate clustering performance.

For three-class LCM, we set class prevalences $\boldsymbol{\pi} = (0.6, 0.2, 0.1)$ with the three dinstinct response probability profiles $\underbrace{(0.9, \ldots, 0.9)}_{L}$, $\underbrace{(0.5, \ldots, 0.5)}_{L}$, $\underbrace{(0.1, \ldots, 0.1)}_{L}$. For six-class LCM, we set $\boldsymbol{\pi} = (0.2, 0.2, 0.2, 0.2, 0.1, 0.1)$ with distinct response probability profiles $\underbrace{(0.9, \ldots, 0.9)}_{L}$, $\underbrace{(0.7, \ldots, 0.7)}_{L}$, $\underbrace{(0.5, \ldots, 0.5)}_{L}$, $\underbrace{(0.3, \ldots, 0.3)}_{L}$, $\underbrace{(0.1, \ldots, 0.1)}_{L}$, $\underbrace{(0.01, \ldots, 0.01)}_{L}$. We simulated $R = 60$ data replications under sample size $N = 50, 100$, dimension $L = 50, 100$ under all settings.

#### A5.2 Ajusted Rand Index (aRI)

We use adjusted Rand index (aRI, Hubert and Arabie, 1985) to assess the agreement between two clustering allocations, e.g,. the estimated and the true clusters. aRI is defined by

$$\mathsf{aRI}(\mathcal{C}, \mathcal{C}') = \frac{\sum_{r,c} \binom{n_{rc}}{2} - \left[\sum_r \binom{n_{r\cdot}}{2} \sum_c \binom{n_{\cdot c}}{2}\right] / \binom{N}{2}}{0.5 \left[\sum_r \binom{n_{r\cdot}}{2} + \sum_c \binom{n_{\cdot c}}{2}\right] - \left[\sum_r \binom{n_{r\cdot}}{2} \sum_c \binom{n_{\cdot c}}{2}\right] / \binom{N}{2}},$$

where $n_{rc}$ represents the number of observations placed in the $r$th cluster of the first partition $\mathcal{C}$ and in the $c$th cluster of the second partition $\mathcal{C}'$, $\sum_{r,c} \binom{n_{rc}}{2} (\leq 0.5 \left[\sum_r \binom{n_{r\cdot}}{2} + \sum_c \binom{n_{\cdot c}}{2}\right])$ is the number of observation pairs placed in the same cluster in both partitions and $\sum_r \binom{n_{r\cdot}}{2}$ and $\sum_c \binom{n_{\cdot c}}{2}$ calculates the number of pairs placed in the same cluster for the first and the same cluster for second partition, respectively. aRI is bounded between $-1$ and 1 and corrects for chance agreement. It equals one for identical clusterings and is on average zero for two random partitions; larger values indicate better agreements.

#### A5.3 Additional Remarks about Simulation 2

We remark on the performance of the other three methods. Over all parameter settings investigated here, the traditional LCA performed worst in the recovery of true clusters (aRI $< 0.68$). The advantage of RLCM comes from the regularization of estimated response probability profiles towards a scientific structure that improves finite-sample clustering performance. The likelihood function of subset clustering is a special case of the RLCM that assumes a non-parsimonious $Q = I_L$ and therefore loses power for detecting clusters compared to RLCM that estimates a structured $Q$ with multiple non-zero elements in its rows. HC is fast and recovers the true clusters reasonably well (ranked second or first among the four methods for more than two thirds of the parameter settings here). The performance of HC is particularly good under a low level of measurement errors ($\theta_0^- = 0.05$) and a large number of relevant features per machine and sometimes performs much better than traditional LCA and subset clustering (e.g., $L = 200$, $N = 50$, $\theta_\ell^+ = 0.8$, $\theta_\ell^- = 0.05$ in Figure S1). The HC studied here requires a pre-specified number of clusters to cut the dendrogram at an appropriate level and produces clusters that require separate methods for uncertainty assessment (e.g., Suzuki and Shimodaira, 2006). The proposed Bayesian RLCM, in contrast, enjoys superior clustering performance and provides direct internal assessment of the uncertainty of clusters and measurement error parameters through the posterior distribution.

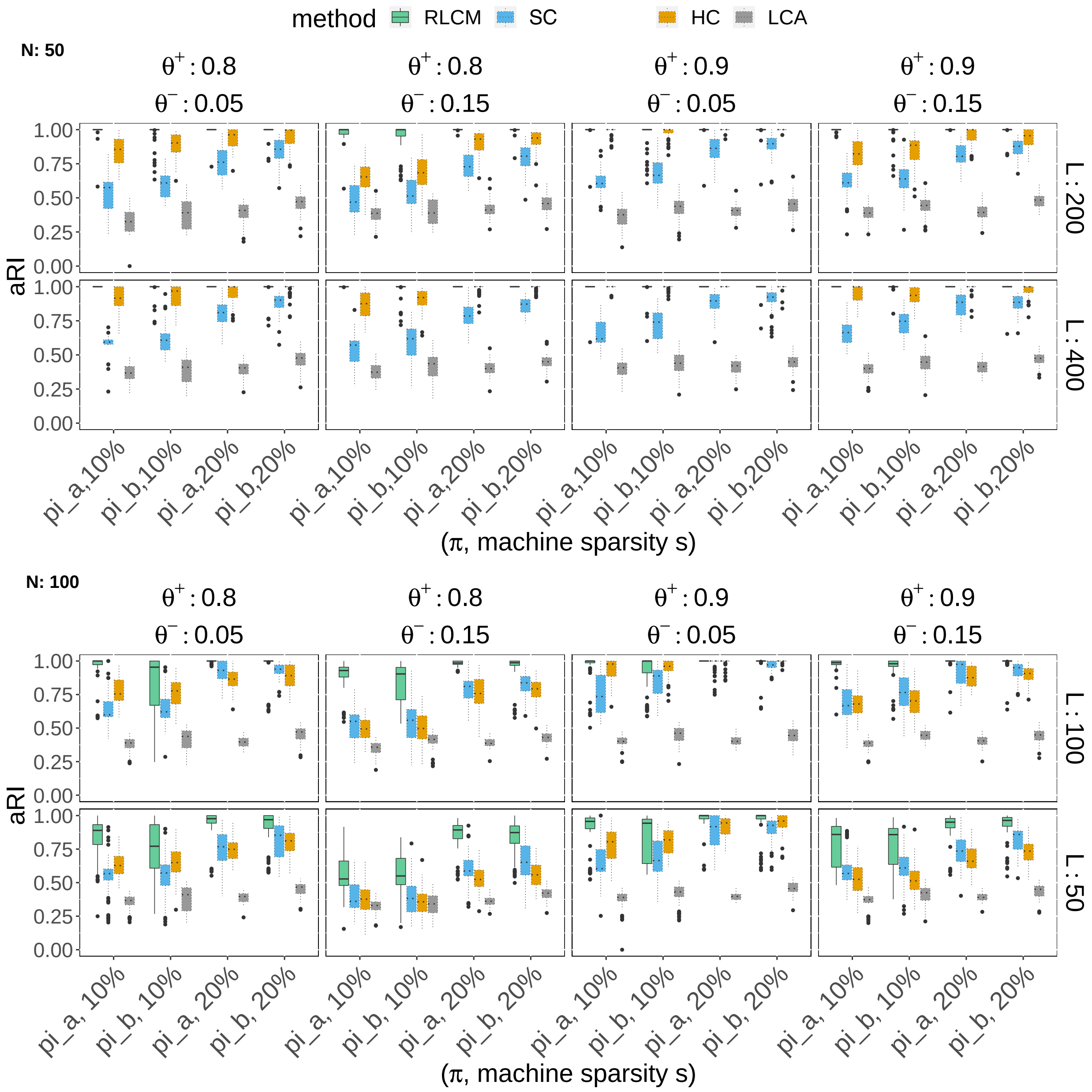


Figure S1: Similar to Figure 3 in the Main Paper, the proposed Bayesian RLCM shows better finite-sample clustering performance than three alternatives over the parameter settings in Simulation 2.

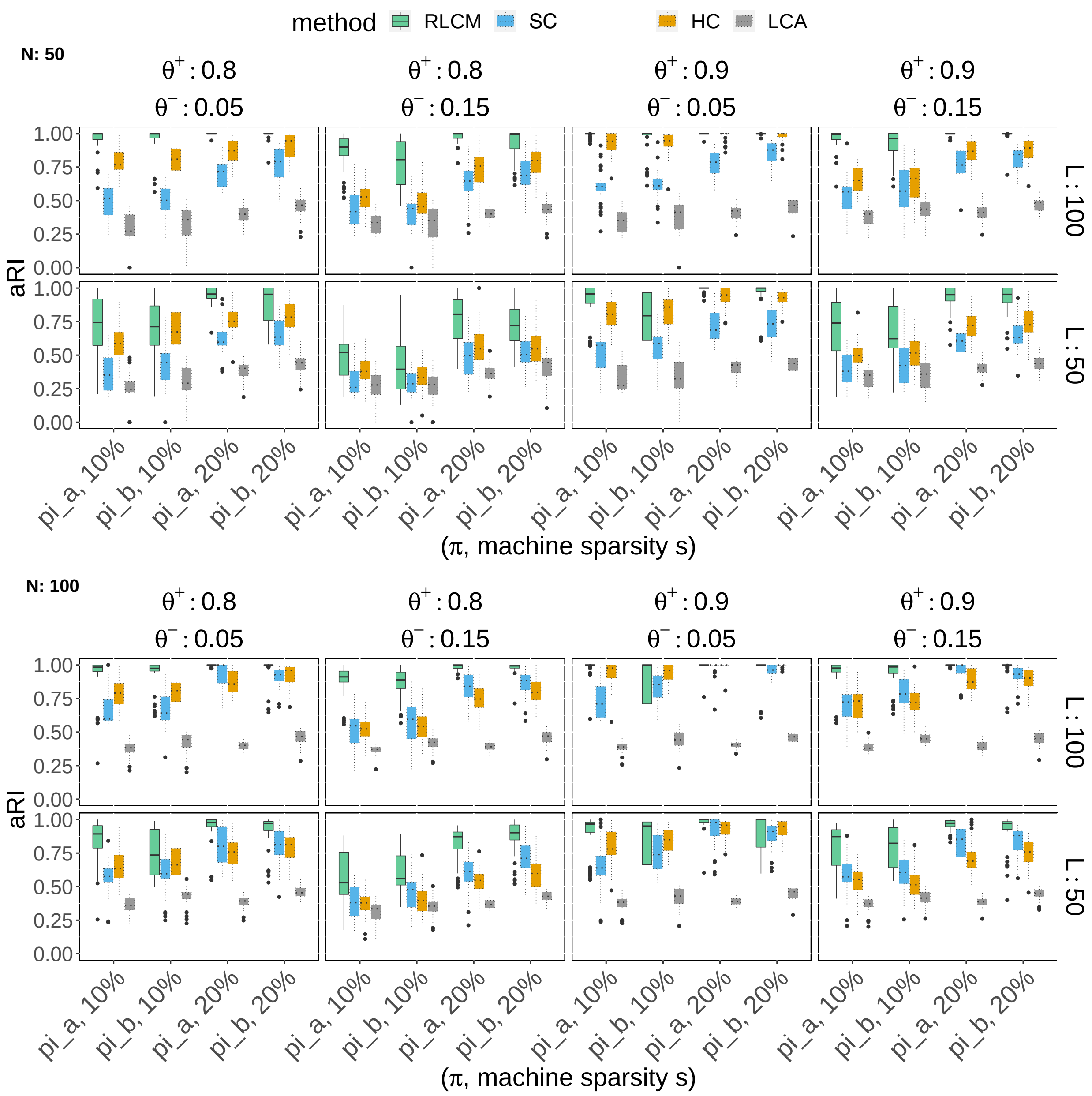


Figure S2: Comparison of clustering performance under the first set of data generating mechanisms (DGM) that have small degrees of deviations from the main model assumptions.

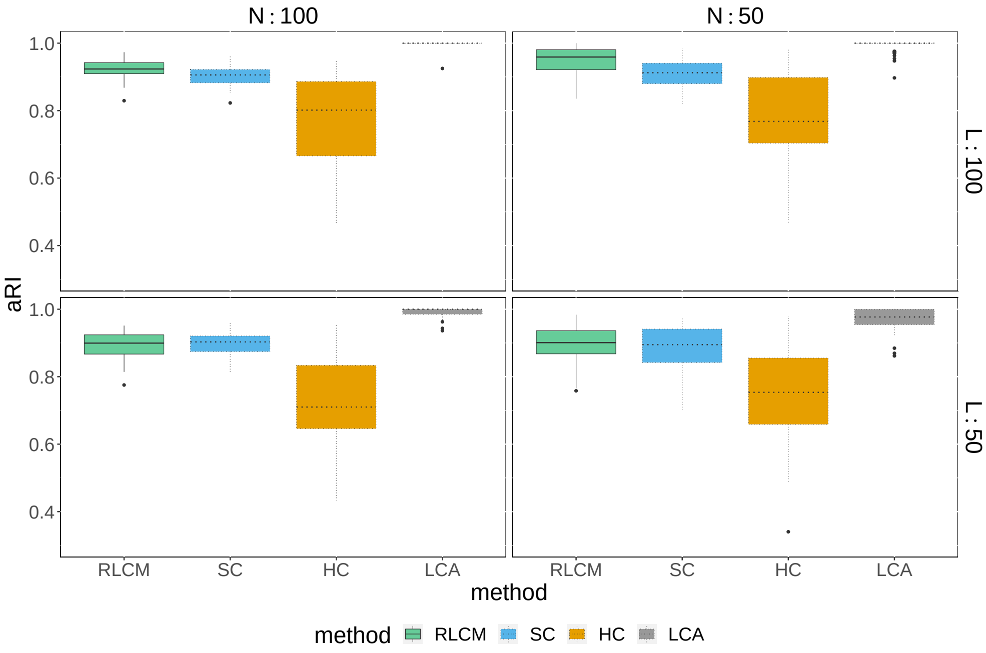


(a) True DGM: three-class LCM

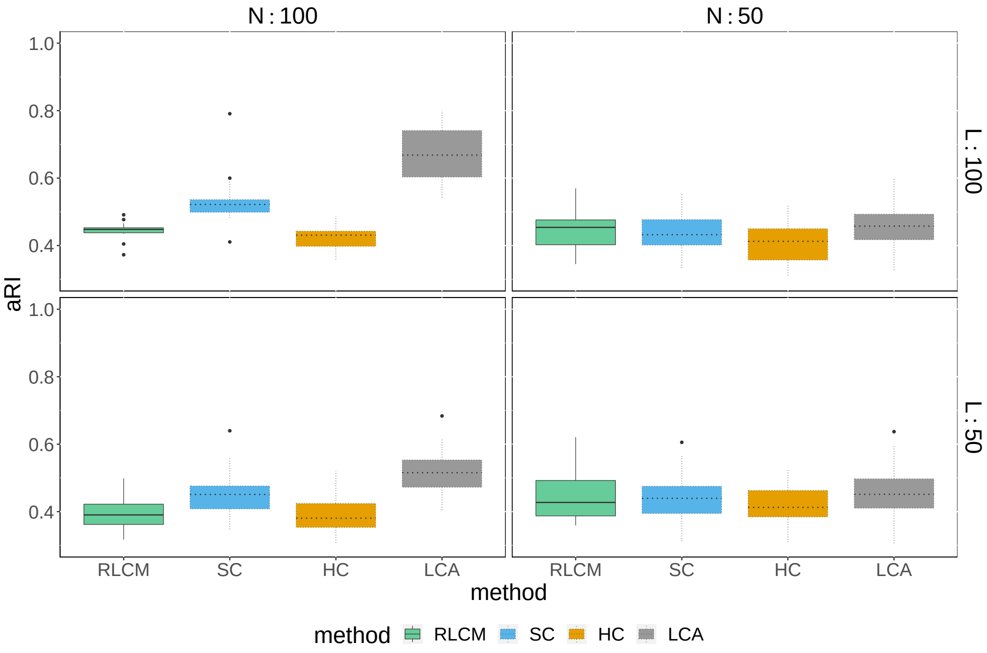


(b) True DGM: six-class LCM

Figure S3: Comparison of clustering performance under a second set of data generating mechanisms (DGMs) that have large degrees of deviations from the assumed model: general LCMs without structural restrictions (a: three-class LCM; b: six-class LCM; See A5.1 for specific parameter values). Because RLCM is not flexible enough to capture the present data generating mechanism, the clustering performance of RLCM is less competitive.

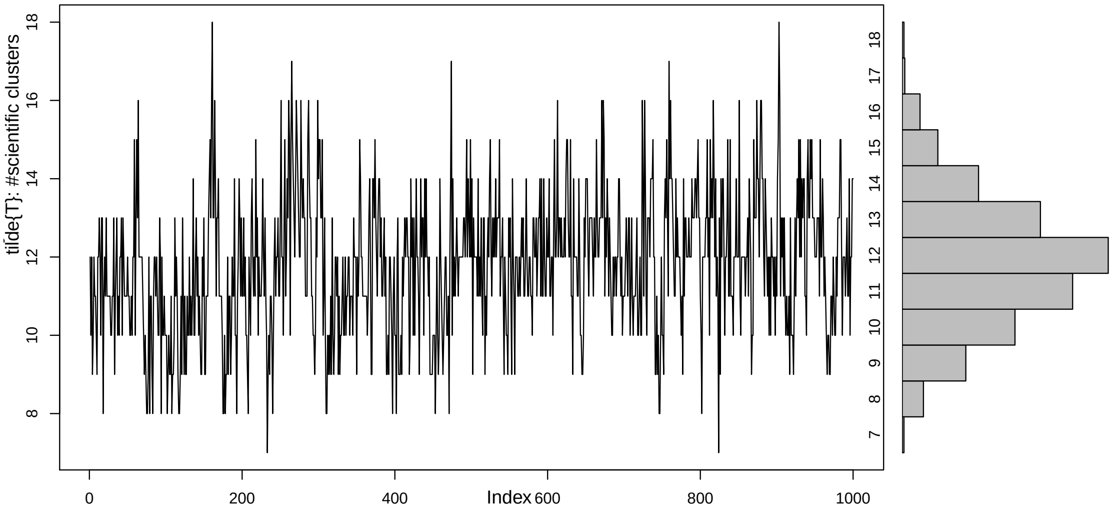


Figure S4: MCMC samples of the number of scientific clusters ($\widetilde{C}$) with its marginal posterior on the right margin.

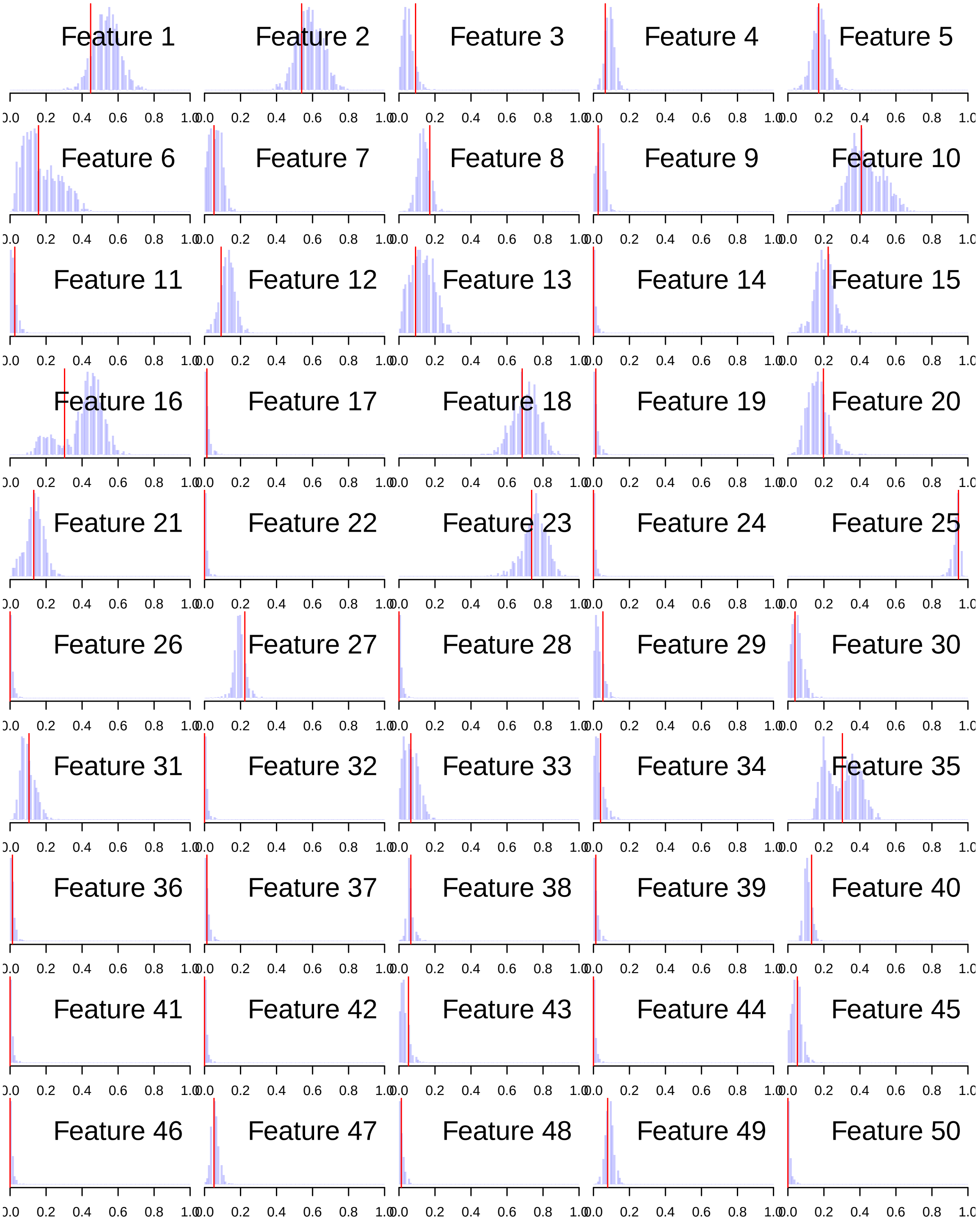


Figure S5: Observed marginal positive rate (solid vertical line) plotted against the posterior predictive distributions for $L = 50$ landmarks (Section 4.2 in the Main Paper).

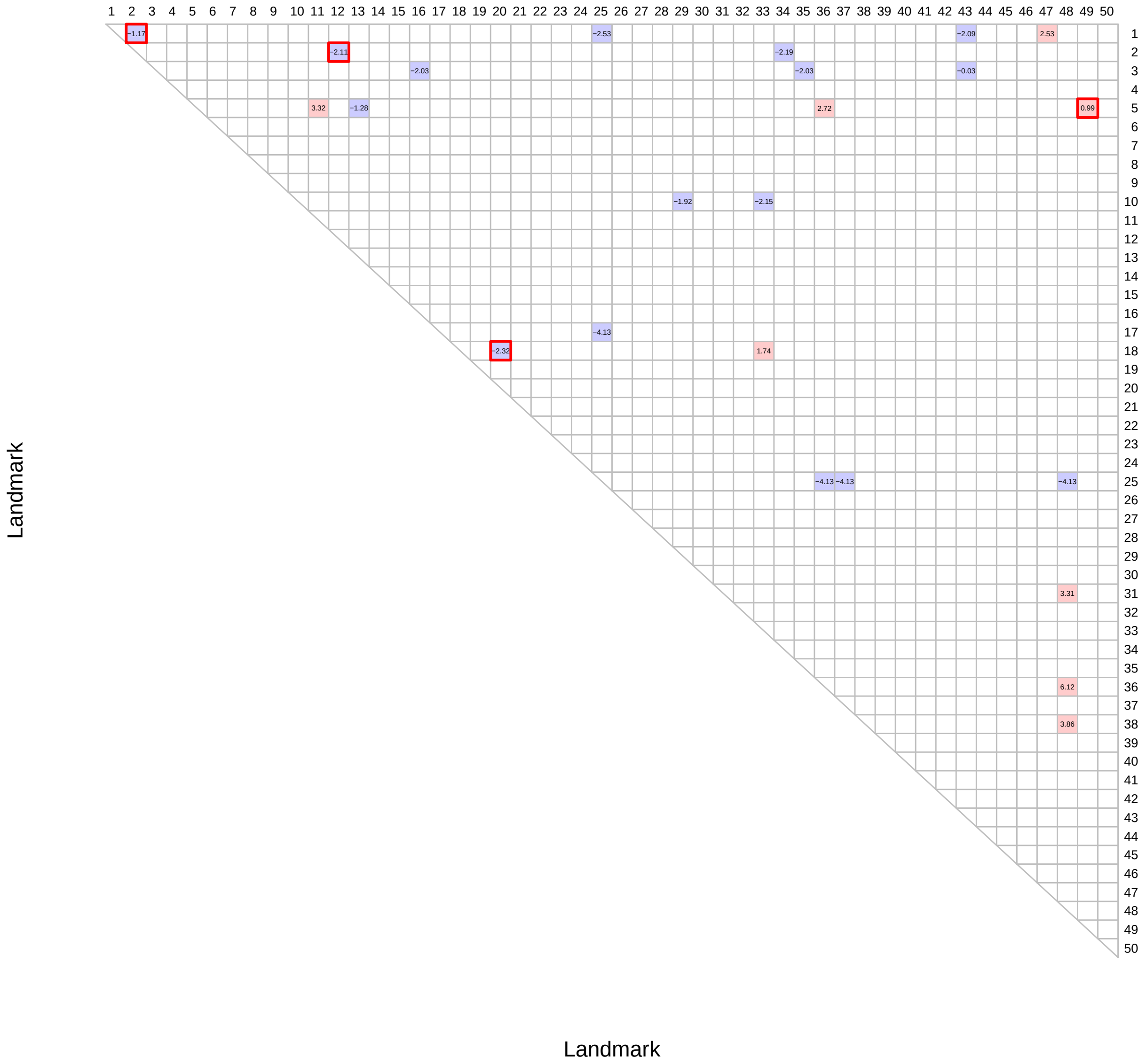


Figure S6: Significant deviations of model predicted log odds ratios (LOR) from the observed LOR. A blank cell indicates a good model prediction for the observed pairwise LOR ($|\mathsf{SLORD}| < 2$); A red (blue) cell indicates model under- (over-) fitting $\mathsf{SLORD} > 2 (< -2)$, where standardized LOR difference (SLORD) is defined as the observed LOR for a pair of landmarks minus the mean LOR for the predictive distribution value divided by the standard deviation of the LOR predictive distribution. A red box indicate that the pair of landmarks have cell counts in the 2 by 2 observed marginal table all greater than 5.

### References

Brooks, S. and Gelman, A. (1998). General methods for monitoring convergence of iterative simulations. *Journal of Computational and Graphical Statistics*, 7(4):434–455.

Geweke, J. and Zhou, G. (1996). Measuring the pricing error of the arbitrage pricing theory. *The review of financial studies*, 9(2):557–587.

Hubert, L. and Arabie, P. (1985). Comparing partitions. *Journal of classification*, 2(1):193–218.

Jain, S. and Neal, R. M. (2004). A split-merge markov chain monte carlo procedure for the dirichlet process mixture model. *Journal of Computational and Graphical Statistics*, 13(1):158–182.

Miller, J. W. and Harrison, M. T. (2017). Mixture models with a prior on the number of components. *Journal of the American Statistical Association*, pages 1–17.

Suzuki, R. and Shimodaira, H. (2006). Pvclust: an r package for assessing the uncertainty in hierarchical clustering. *Bioinformatics*, 22(12):1540–1542.
